# *Il33* expressing cDC2s promote expansion of ILC2s and eosinophilia in fungal airway inflammation in male mice

**DOI:** 10.64898/2025.12.02.691763

**Authors:** Lisa-Marie Graf, Daniel Radtke, Andreas Ruhl, Kirstin Castiglione, Stefan Wirtz, Sven Krappmann, Barbara U. Schraml, David Voehringer

## Abstract

Eosinophilic allergic asthma is often associated with fungal sensitization and represents the dominant form of asthma in adolescents. The mechanisms by which lung eosinophilia is regulated in this context are incompletely understood. Here, we demonstrate that type 2 innate lymphoid cells (ILC2s) are generally required in addition to Th2 cells for *Aspergillus fumigatus*-elicited eosinophilic lung inflammation in mice and this effect was independent of ILC2-derived IL-5. Surprisingly, expansion of ILC2s in male mice was ST2-independent, but required expression of *Il33* in cDC2s induced by Th2-derived IL-4/13. This IL-33-plus IL-4-regulated transcriptional module in cDC2s included expression of factors associated with activation of ILC2s and was restricted to CCR7^+^ cDC2s in the inflamed lung. Our findings uncovered a novel intimate cross-talk between IL-33^+^ cDC2s, ILC2s and Th2 cells to orchestrate eosinophilia in a sex-dependent manner upon *A. fumigatus*-induced allergic lung inflammation.

## Introduction

Asthma is a heterogenous chronic disease of the lung which currently affects about 260 million people worldwide and causes a major burden on patients, health care systems and economies (*1*). Asthma can be caused by many factors including sensitization to environmental fungi such as *A. fumigatus* (*Af*). Inhalation of *Af* conidia (spores) may result in severe asthma with fungal sensitization (SAFS) or allergic bronchopulmonary aspergillosis (ABPA), an eosinophilic inflammatory response found mainly in asthmatic or cystic fibrosis patients (*2*). Both conditions are characterized by a strong type 2 immune response including pronounced lung eosinophilia (*3*). However, underlying immunological mechanisms are poorly understood (*4*). In the past, we reported that type 2 T helper (Th2) cells are critical for induction of lung eosinophilia in a murine model of *Af*-elicited lung inflammation (*5*).

Besides Th2 cells, type 2 innate lymphoid cells (ILC2s) have been shown to play a crucial role in allergic airway inflammation (*6*). They are present in low numbers at steady-state conditions in the lung. However, allergen-derived proteases and other factors can activate or damage lung epithelial cells leading to release of alarmins (IL-25, IL-33, TSLP), which are potent activators of ILC2s (*7*). Activated ILC2s then secrete IL-5, IL-9, IL-13, and to lesser extent IL-4 to promote a type 2 immune response (*8*). ILC2s were found to be the most critical cellular source of IL-5 which plays an important role for development and survival of eosinophils and B1 cells (*9, 10*). ILC2-derived IL-9 and IL-13 have further been shown to act on conventional dendritic cells (cDCs) and elicit their migration into draining lymph nodes and differentiation into Th2-polarizing cDC2s (*11, 12*). Therefore, the absence of ILC2s not only alleviates lung eosinophilia but generally reduces asthmatic symptoms in different models of allergic airway inflammation (*13, 14*). However, it has not been investigated whether ILC2s are critical players in *Af*-induced allergic lung inflammation (*15*).

Interestingly, asthma prevalence in children is higher in the male as compared to the female population and this bias is reversed after puberty (*1*). Studies in mice have shown that androgen signaling in ILC2s dampens their development and activation in allergen-challenged male mice (*16, 17*). This sex disparity in ILC2s is reflected in a greater proportion of type 2 cytokines and eosinophils in the lung of female as compared to male mice after allergen challenge (*18*).

We hypothesized that ILC2s are important players in *Af*-induced lung eosinophilia and aimed at dissecting the role of cDC2s, Th2 cells and ILC2s as well as potential sex-dependent differences in this response. Indeed, we found that beside Th2 cells, ILC2s are required for induction of eosinophilia but not for accumulation of Th2 cells in the lung. Surprisingly, this effect was not mediated by ILC2-derived IL-5. In male mice, IL-33-expressing cDC2s and Th2-derived IL-4/IL-13 were required for expansion of the ILC2 population in the lung. IL-33 expression in DCs could be induced by IL-4 in a STAT6-dependent manner. Further, an IL-33-dependent and IL-4-induced gene expression profile including activating signals for ILC2s was found in migratory CCR7^+^ cDC2s in the lung of *Af*-challenged mice. These results indicate that Th2 cells feedback on cDC2s, which are crucial for expansion of ILC2s in the lung of male mice probably by interfering with inhibitory signals downstream of the androgen receptor. Overall, our study provides new insights in sex-dependent regulation of *Af*-induced lung eosinophilia via a novel cross-talk between IL-33 expressing cDC2s, Th2 cells and ILC2s.

## Results

### Type 2 innate lymphoid cells are essential for the induction of eosinophilia in *Af*-induced lung inflammation

In the past we and others reported that repeated intranasal treatment with a low number of live *Af* conidia induces a strong Th2 response and eosinophilia (*5, 19*). To investigate the contribution of ILC2s to *Af*-induced lung inflammation, we treated mice four times intranasally with 2×10^6^ live conidia (Fig. 1A) and examined the expansion of the ILC2 population in the lung. ILC2s were significantly increased in the early phase of inflammation (up to day 14) but then declined again, thus *Af*-elicited immunity was analyzed at day 14 (Fig. S1A, B). As sex-differences play a role for ILC2 expansion in asthma, we analyzed both female and male mice. ILC2 numbers were significantly increased in males, and a similar trend was observed in females (Fig. 1B). While IL-13^+^ ILC2s were significantly increased in mice of both sexes (Fig. S1D), total IL-5^+^ ILC2s were only increased in the lungs of male mice, as females already displayed high basal levels in saline treated mice (Fig. S1E). Furthermore, the increase of Th2 cells and eosinophils was comparable between males and females (Fig. 1C, D). As it has been proposed that protease-mediated tissue damage promotes expansion and activation of ILC2s in asthma, we investigated whether *Af*-derived proteases contribute to this response. We treated mice with an *Af* strain which lacks extracellular protease activity (*Af* prtTΔ) but did not observe significant differences in ILC2, Th2, and eosinophil numbers in the lung (Fig. S1F-H).

**Figure 1:**
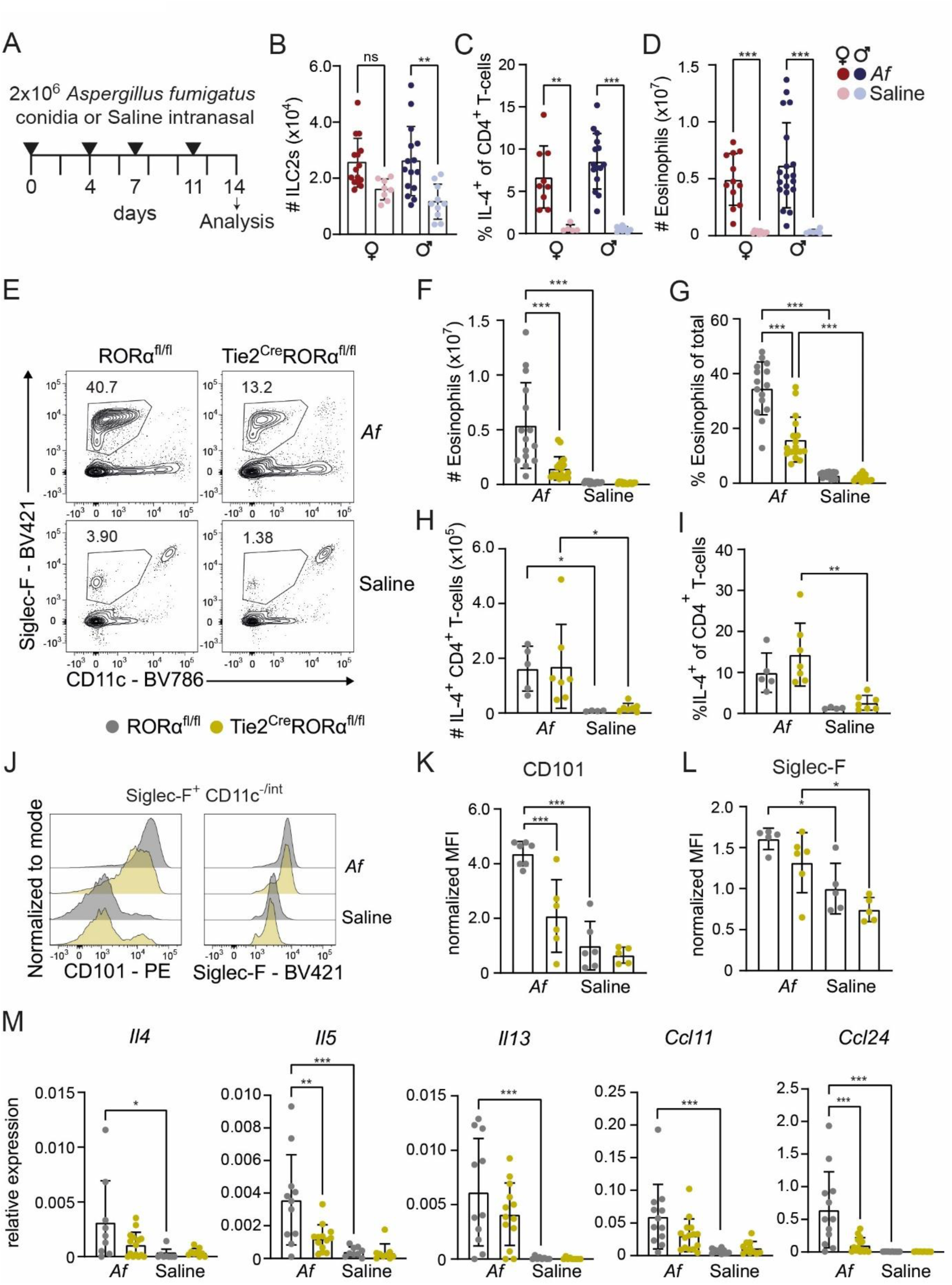
Repeated intranasal administration of low dose *Aspergillus fumigatus* (*Af*) conidia elicits ILC2-dependent lung eosinophilia. (A) Treatment scheme used to induce *Af-*elicited allergic lung inflammation with analysis on day 14. (B-D) Total numbers of ILC2s (B), percentage of IL-4^+^ T-cells (C) and total numbers of eosinophils (D) separated by sex in the lung of wild-type (WT) C57BL/6 mice. (E) Flow cytometry plots show gating of eosinophils (Siglec-F^+^CD11c^−/lo^) in the lung of RORα^fl/fl^ and Tie2^Cre^RORα^fl/fl^ mice. (F-I) Total numbers (F) and percentage (G) of eosinophils in the lung as well as total numbers (H) and percentages (I) of IL-4^+^CD4^+^ T cells. (J) Histograms show expression levels of CD101 and Siglec-F on eosinophils (Siglec-F^+^CD11c^−/int^) in the lung. (K, L) Fluorescence intensity of CD101 (K) and Siglec-F (L) normalized to the mean fluorescence intensity of saline treated WT mice. (M) Quantitative RT-PCR from total lung tissue normalized to *Hprt*. Bar graphs show mean±SEM for n≥5 (B-D), n≥14 (F, G), n≥5 (H, I, K, L), n≥10 (M) mice per group pooled from 7 (B-D), 5 (F, G), 2 (H, I, K, L) or 4 (M) independent experiments. One-way ANOVA was used for statistical analysis (* p<0.05, ** p<0.01, *** p<0.001). Only relevant statistical significances are indicated.

Next, we determined whether ILC2s are relevant for the inflammatory response and therefore used an ILC2-deficient mouse strain (Tie2^Cre^RORa^fl/fl^, Fig. S1I, J). Since an impact of ILC2s on Th2 induction has been shown in some allergic airway inflammation models (*12, 13*), we also examined the Th2 response but did not observe reduced Th2 cells in ILC2-deficient animals (Fig. 1H, I, Fig. S1L). Eosinophilia, as a hallmark of *Af*-induced lung inflammation, was significantly reduced in mice lacking ILC2s (Fig.1 E-G), and this effect was similar for both sexes (Fig. S1K). Moreover, eosinophils from ILC2-deficient animals showed a reduced inflammatory phenotype indicated by reduced levels of CD101 expression (Fig. 1J, K). In line with reduced eosinophilia, expression of *Il5* and *Ccl24* encoding the eosinophil-recruiting chemokine CCL24 (also known as eotaxin-2) was significantly reduced in whole lung tissue, with a similar trend for *Il4*, *Il13* and *Ccl11* (Fig. 1M).

This data indicates that ILC2s prominently contribute to eosinophilia and to the inflammatory profile of eosinophils in *Af*-induced lung inflammation independently of Th2 cells.

### ILC2-derived IL-4/IL-13 promotes eosinophil recruitment only in male mice

To better understand the contribution of ILC2s to eosinophilia, we investigated the role of ILC2-derived cytokines. First, we addressed the role of IL-5 as ILC2s have been shown to be the main source of this cytokines and IL-5 is critical for induction of eosinophilia(*9, 20*). Therefore, we generated mixed bone marrow chimeras by lethal irradiation of ILC2-deficient hosts and reconstitution with a mixture of 20% IL-5-deficient (Red5 mice) or wild-type (WT) and 80% ILC2-deficient bone marrow (Fig. 2A). By this approach we generated mice in which either all ILC2s could produce IL-5 (ILC2^WT^) or were deficient in IL-5 production (ILC2^IL-5k.o.^). Apart from IL-5^+^ ILC2s (Fig. 2B, C) total ILC2s, IL-13^+^ ILC2s and IL-5^+^ Th2 cells could normally develop in ILC2^IL-5k.o.^ mice (Fig. S2B-D). Unexpectedly, lung eosinophilia was comparable between ILC2^IL-5k.o.^ and ILC2^WT^ animals (Fig. 2D, E), and independent of their sex (Fig. S2E).

**Figure 2:**
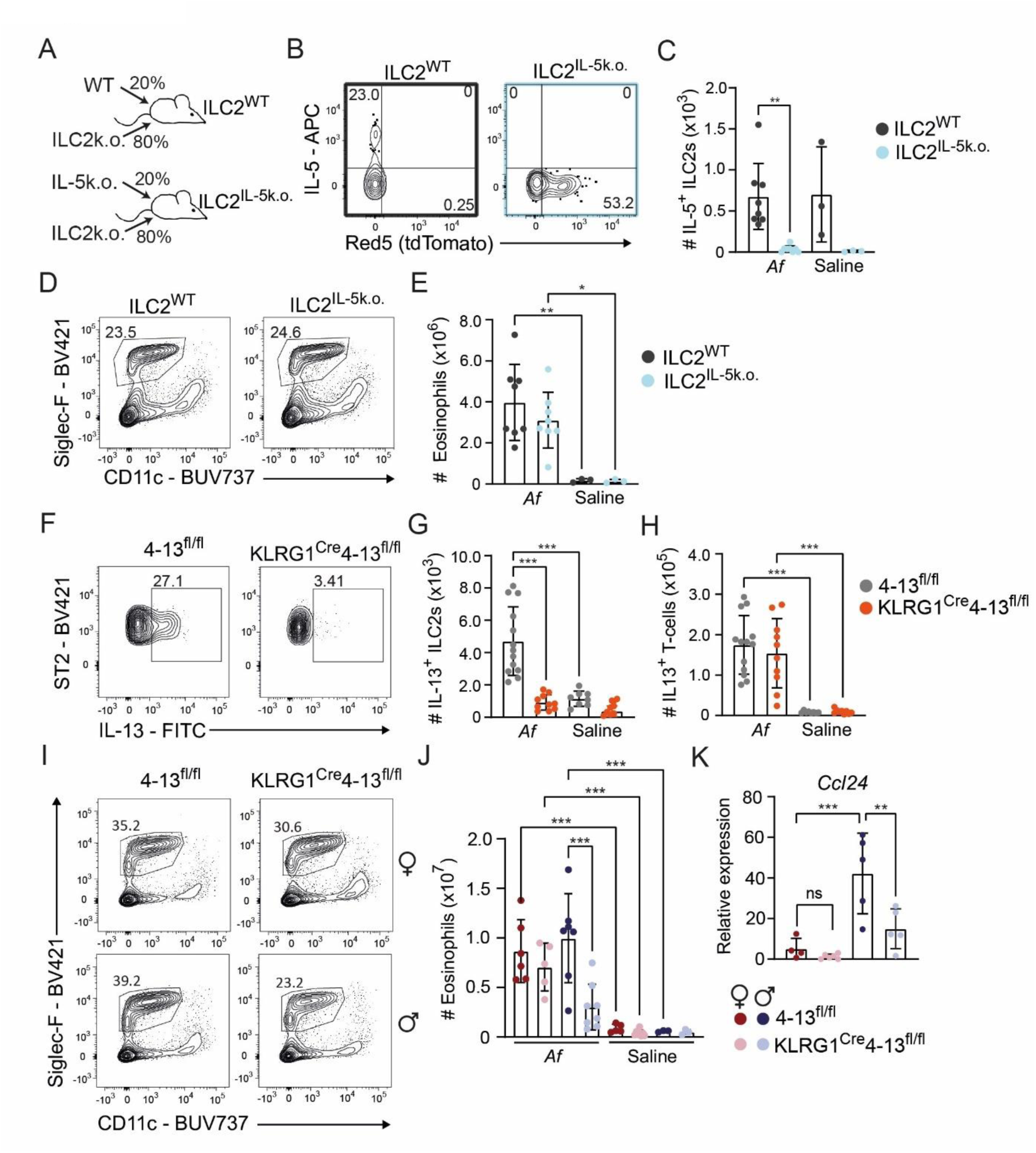
Sex bias in ILC2-mediated eosinophilia in *Af-*induced lung inflammation. (A) Lethally irradiated ILC2-deficient mice (Tie2^Cre^RORα^fl/fl^) were reconstituted with 20% WT or 20% IL-5k.o. (Red5) and 80% ILC2k.o. (Tie2^Cre^RORα^fl/fl^) bone marrow (BM). (B) Flow cytometry plots show representative staining of intracellular IL-5 and Red5 (tdTomato) signal in lung ILC2s (gated to viable Lin^−^CD45^+^ST2^+^CD127^+^ cells) from chimeras reconstituted with WT or IL-5k.o. BM. (C) Total numbers of IL-5^+^ ILC2s. (D) Flow cytometry plots show frequencies of gated eosinophils (Siglec-F^+^CD11c^−/int^. (E) Quantification of eosinophils in D. (F) Representative flow cytometry plots of intracellular IL-13 staining in ILC2s from KLRG1^Cre^4-13^fl/fl^ and 4-13^fl/fl^ control mice (gated to viable Lin^−^CD45^+^ST2^+^CD127^+^ cells). (G, H) Total numbers of IL-13^+^ ILC2s (G) and IL-13^+^ T cells (H). (I) Flow cytometry plots show frequencies of gated eosinophils (Siglec-F^+^ CD11c^−/int^) in male and female mice. (J) Quantification of eosinophils from (I). (K) Sorted lung macrophages (gated as CD64^+^MerTK^+^ cells, Fig. S2H) were analyzed by quantitative RT-PCR. Bars display relative expression of *Ccl24* normalized to *Hprt*. Graphs show mean±SEM for n=3-8 chimeras (C, E), n=3-8 mice (G, H, J), n≥4 mice (K) per group pooled from two (C, E), four (G, H), five (J) or two (K) independent experiments. One-way ANOVA was performed for statistical testing (* p<0.05, ** p<0.01, *** p<0.001). Only relevant statistical significances are indicated.

This result raised the possibility that ILC2-derived IL-4 and IL-13 rather than IL-5 is important for induction of lung eosinophilia, e.g. by inducing expression of eosinophil-recruiting chemokines. Therefore, we generated mice that lack IL-4/IL-13 expression in ILC2s by crossing KLRG1^Cre^tdTomato^stop-flox-stop^ mice to conditional IL-4/IL-13-deficient (4-13^fl/fl^) mice. Although only a fraction of ILC2s was positive for KLRG1 expression on the cell surface, almost all lung ILC2s had expressed KLRG1 at some point during their development as indicated by the tdTomato fate-mapping signal (Fig. S2F, G). Flow cytometric analysis of intracellular IL-13 confirmed that ILC2s from KLRG1^Cre^4-13^fl/fl^ mice lack IL-13 expression, whereas IL-13 expression in Th2 cells is not affected (Fig. 2F-H). After intranasal treatment of this mouse strain with *Af* conidia, sex-specific differences in lung eosinophilia were observed. Male mice displayed a significantly reduced number of eosinophils, when lacking IL-4/IL-13 in ILC2s, while this was not the case for female mice (Fig. 2I-J).

To further investigate this sex-dependent difference of eosinophilia in KLRG1^Cre^4-13^fl/fl^ mice, we considered differential expression of eotaxins as a potential explanation. As macrophages are a major source of CCL24 and its expression can be induced by IL-4/IL-13 signaling (*21*), we analyzed macrophages sorted from the lung of *Af*-challenged male and female KLRG1^Cre^4-13^fl/fl^ mice. Surprisingly, in control mice, male-derived macrophages expressed significantly higher levels of *Ccl24* mRNA than female-derived macrophages, and this effect was largely dependent on IL-4/IL-13 from ILC2s (Fig. 2K). In contrast, *Ccl11* was expressed at a similar level in whole lung tissue of control mice of both sexes, and non-significantly reduced in female mice lacking IL-4/IL-13 in ILC2s (Fig. S2I).

Overall, these results indicate that ILC2-derived IL-4/IL-13 plays an important non-redundant role in eosinophil recruitment to the lung of *Af*-challenged male mice most likely by promoting CCL24 expression in lung macrophages.

### cDC2s and Th2 cells promote expansion of ILC2s in the lung of *Af*-challenged male mice

Next, we sought to investigate whether cDC2s and Th2 cells, as other key players of type 2 immunity, are involved in male-specific regulation of lung eosinophilia. Therefore, two different mouse strains were used in which the transcription factor IRF4, required for differentiation of Mgl2^+^ cDC2s, is deleted either during DC development (Clec9a^Cre^Irf4^fl/fl^ mice) (*22*) or in CD11c-expressing cells including cDCs (CD11c^Cre^Irf4^fl/fl^ mice). In agreement with previous studies (*23, 24*), male and female mice lacking *Irf4* in DCs did neither develop a decent Th2 response nor eosinophilia (Fig. 3A, B). However, the number of ILC2s was significantly reduced only in male cDC2-deficient mice as compared to control mice (Fig. 3C, D).

**Figure 3:**
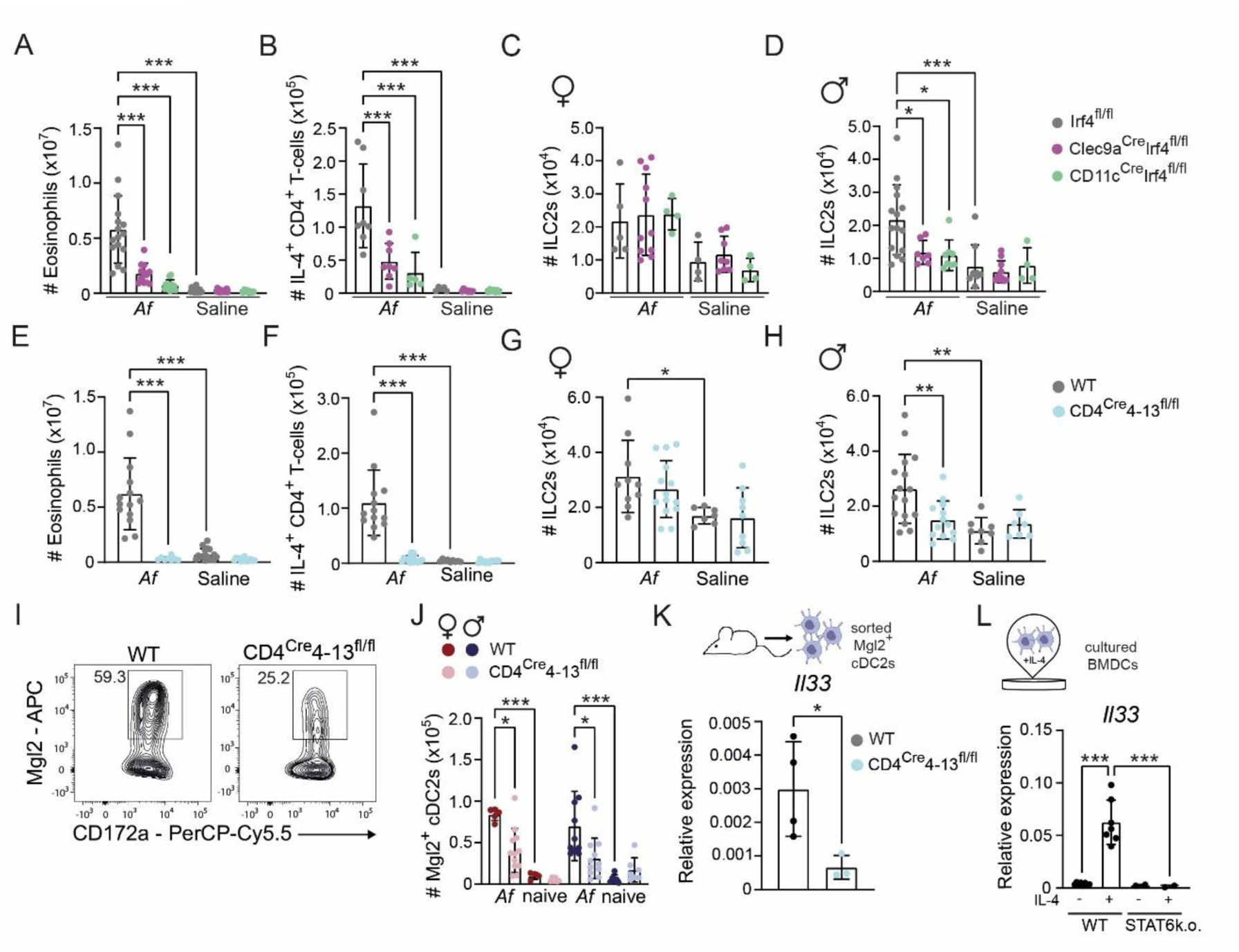
Crosstalk between cDC2s and Th2 cells promotes expansion of ILC2s in the lung of *Af*-challenged male mice. (A-D) Irf4^fl/fl^, Clec9a^Cre^Irf4^fl/fl^ and CD11c^Cre^Irf4^fl/fl^ mice were analyzed on d14 after initial *Af* treatment. Total numbers of eosinophils (A), IL-4^+^CD4^+^ T cells (B) and ILC2s in females (C) and males (D) are displayed. (E-H) CD4^Cre^4-13^fl/fl^ and control mice were analyzed on d14 after initial *Af* treatment. Total numbers of eosinophils (E), IL-4^+^CD4^+^ T cells (F) and ILC2s in females (G) and males (H) were determined. (I) Flow cytometry plots show gating of Mgl2^+^ cDC2s (gated on viable Siglec-F^−^CD64^−^MerTK^−^MHC-II^+^CD11c^+^CD11b^+^ cells, Fig. S3A). (J) Quantification of (I) (Mgl2^+^ cDC2s). (K) Mgl2^+^ cDC2s were sorted by flow cytometry from lungs of CD4^Cre^413^fl/fl^ or control mice on d14 after initial *Af* treatment. Quantitative RT-PCR of *Il33* normalized to *Hprt* of sorted Mgl2^+^ cDC2s (Gating strategy Fig. S3B). (L) Quantitative RT-PCR of *Il33* normalized to *Hprt* from unstimulated or IL-4 (5 ng/ml) stimulated BMDCs cultured from WT or STAT6-deficient (STAT6k.o.) mice from 3 (WT) or 2 (STAT6k.o.) independent experiments. Bar graphs display mean±SEM from n=5-17 (A-D), n=7-16 (E-H), n=4-12 (J), n=3-4 (K) mice pooled from 7 (A-D), 5 (E-H, J), and 3 (K) independent experiments. Statistical analysis was performed using one-way ANOVA (A-H, J, L) and unpaired Student’s t-test (K) (* p<0.05, ** p<0.01, ***p<0.001). Only relevant statistical significances are indicated.

As Th2 cells have been described to promote expansion of ILC2s in some experimental settings (*13, 25*), we determined the ILC2 response in mice lacking IL-4/IL-13 producing T cells (CD4^Cre^4-13^fl/fl^). While T cell-derived IL-4/IL-13 was essential for lung eosinophilia in both male and female mice, only male mice showed significantly reduced expansion of ILC2s in CD4^Cre^4-13^fl/fl^ mice upon *Af*-induced lung inflammation (Fig. 3E-H).

We considered the possibility that T cell-derived IL-4/IL-13 acts on cDC2s to promote expansion of ILC2s in male mice. It has been reported that cDC2s stimulated with house dust mite extract express *Il33* a potent activator of ILC2s (*24*). Moreover, an *in silico* prediction tool indicated that the IL-4/IL-13-activated transcription factor STAT6 is a potential inducer of *Il33* expression (*26*). Therefore, we first determined the total number of Mgl2^+^ cDC2s in the lung of CD4^Cre^4-13^fl/fl^ and control mice (Fig. 3I). In both, females and males, the total number of Mgl2^+^ cDC2s was significantly reduced in mice lacking expression of IL-4/IL-13 from T cells (Fig. 3J). Further Mgl2^+^ cDC2s sorted from the lung of mice lacking IL-4/13 producing T-cells showed significantly lower levels of *Il33* mRNA expression (Fig. 3K). We further confirmed in bone marrow derived DCs (BMDCs) that IL-4-induced expression of *Il33* is STAT6-dependent (Fig. 3L).

These results suggest a crosstalk between cDC2s and Th2 cells which is important for induction of ILC2s and subsequent lung eosinophilia in male mice.

### IL-33-expressing DCs induce expansion of ILC2s in male animals

After revealing that Th2-derived cytokines can induce *Il33* expression in DCs, we addressed the function of IL-33 in DCs for the type 2 immune response in male mice. To this end, we generated mixed BM chimeras with a mixture of 20% WT or IL-33k.o. and 80% Clec9a^Cre^Irf4^fl/fl^ BM (Fig. 4A). In mice reconstituted with IL-33k.o. BM, only cDC2s deficient in *Il33* will develop (cDC2^IL-33k.o.^), while cDC2s of the control group can express *Il33* (cDC2^WT^). First, we examined the impact of *Il33*-expressing cDC2s on the T cell compartment. For Th2 cells and Foxp3^+^ regulatory T cells no significant difference was observed between cDC2^WT^ and cDC2^IL-33k.o^ mice (Fig. 4B-D). Despite normal induction of Th2 cells, ILC2 numbers and eosinophilia were decreased in cDC2^IL-33k.o.^ mice compared to cDC2^WT^ controls (Fig. 4E, F).

**Figure 4:**
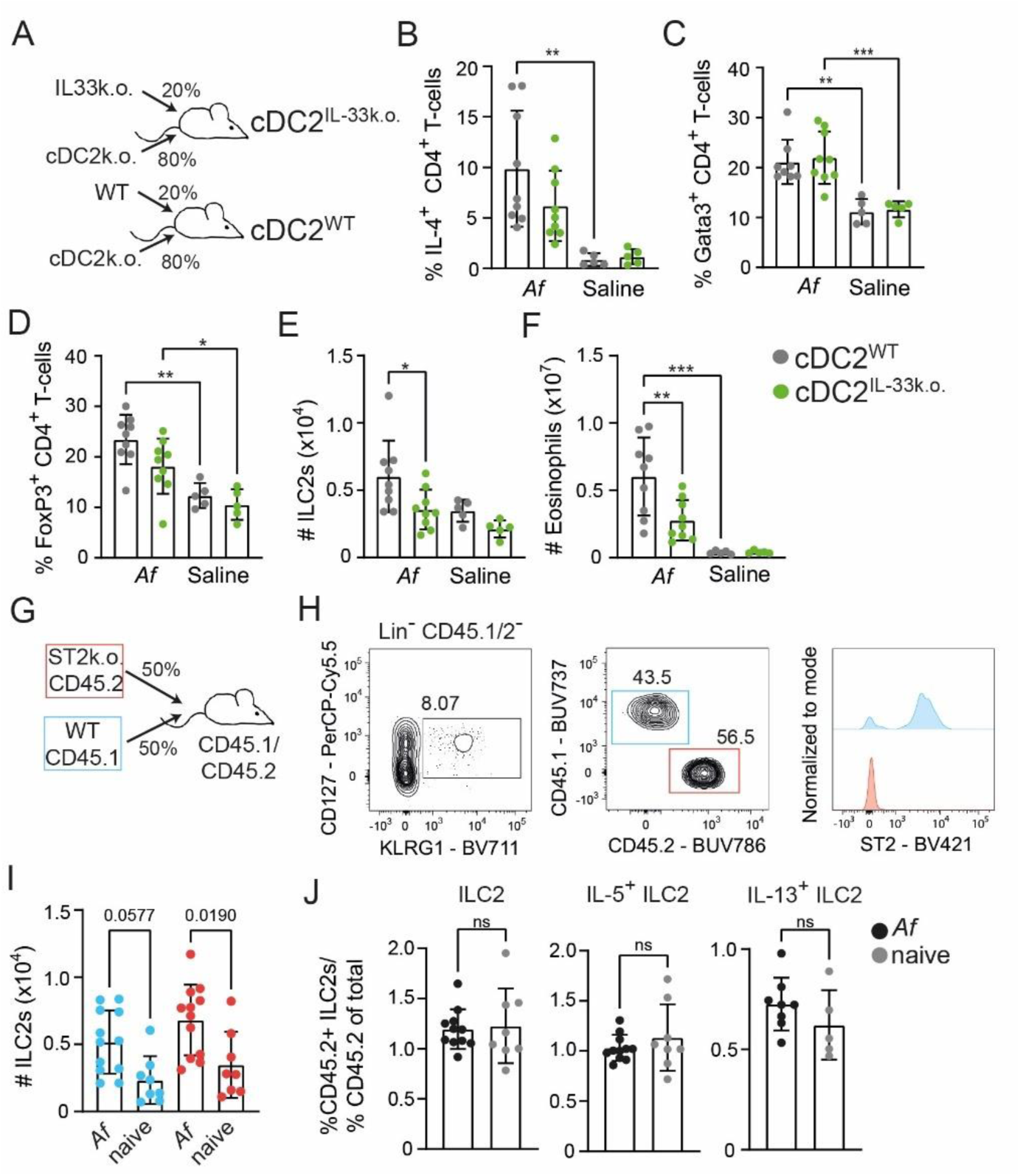
cDC2-intrinsic IL-33 is important for ILC2 expansion and eosinophilia in male mice. (A) Lethally irradiated mice were reconstituted with a mixture of 80% Clec9a^Cre^Irf4^fl/fl^ and either 20% WT or 20% IL-33k.o. bone marrow. The reconstituted mice were subjected to the *Af* model or received intranasally saline. (B-D) Frequencies of IL-4^+^ (B), Gata3^+^ (C) and FoxP3^+^ (D) T cells. (E, F) Total numbers of ILC2s (E) and eosinophils (F). (G) Lethally irradiated CD45.1/CD45.2 mice were reconstituted with a mixture of 50% ST2k.o.-CD45.2 and 50% WT-CD45.1 BM. (H) Representative flow cytometric gating of ILC2s (Lin^−^CD45.1/2^−^CD45.1^+^ or CD45.2^+^), further gating to CD45.1^+^ and CD45.2^+^ ILC2s as well as their normalized expression level of ST2. (I) Quantification of H (middle plot) of CD45.1^+^ (blue) and CD45.2^+^ (red) ILC2s in the lung. (J) Bar graphs display the proportion of percentage of CD45.2 cells in indicated target populations divided by the percentage of CD45.2 cells of total cells. Comparing these ratios between Saline treated mice and the *Af*-group reveals if a population is altered through treatment. The resulting ratio is displayed in the bar graphs. Bar graphs display the mean±SEM of n=5-9 (B-F) and n=5-11 (I-J) mice from 2 (B-F) and 2-3 (I-J) individual experiments. Two-way ANOVA (B-F, I) or unpaired Student’s t-test (J) was used for statistical analysis (* p<0.05, ** p<0.01, ***p<0.001). Only relevant statistical significances are indicated.

To establish, whether cDC2-derived IL-33 could have direct effects on ILC2s, expansion of ST2-deficient ILC2s (lack IL-33 receptor) was compared to WT ILC2s in *Af*-induced lung inflammation. Therefore, lethally irradiated CD45.1/CD45.2 heterozygous male mice were reconstituted with a mixture of 50% ST2k.o. (CD45.2) and 50% congenic WT (CD45.1) BM (both from male mice) to study expansion of ST2-deficient and WT ILC2s in the same environmental setting (Fig. 4G). Reconstitution efficiency and ST2-deficiency of CD45.2^+^ cells was confirmed by flow cytometry (Fig. 4H). Total numbers of WT and ST2k.o. ILC2s were both similarly increased after *Af* treatment (Fig. 4I). To further determine the effect of ST2-deficiency, the percentage of CD45.2^+^ (ST2k.o.) ILC2s was compared to the overall percentage of CD45.2^+^ cells. In the steady-state situation (saline-treated control group), the mean ratio of total ILC2s and IL-5^+^ ILC2s was almost at 1.0 or even higher, indicating no ST2 dependency (Fig. 4J). In contrast, for IL-13^+^ ILC2s the mean ratio was 0.6, indicating that IL-33 directly promotes IL-13^+^ ILC2s at steady-state in the lung. Compared to steady-state, the ratio was unchanged upon *Af* treatment, indicating that IL-33 is dispensable for ILC2 expansion during *Af-*induced lung inflammation.

These results indicate that IL-33^+^ cDC2s promote expansion of ILC2s during *Af*-induced lung inflammation in male mice. However, this effect is largely independent of ST2 on ILC2s and suggests a cell-intrinsic role for IL-33 in cDC2s.

### IL-33 regulates a cDC2-intrinsic transcriptional program including factors that act on ILC2s

Considering the fact that previous publications already provided evidence that IL-33 might change transcriptional activity in a cell-intrinsic manner (*27, 28*), we further aimed at identification of transcriptional differences between WT and IL-33-deficient DCs. Therefore, GM-CSF stimulated bone marrow cells, which give rise to a mixture of macrophages and dendritic cells (further termed as BMDCs) (*29*), derived from WT or IL-33-deficient mice were either left untreated or were stimulated with IL-4 and bulk RNA sequencing analysis was performed.

Principle component analysis shows a major difference in gene expression profiles of WT BMDCs upon IL-4 treatment, driven especially by STAT6-regulated genes such as *Ccl24*, *Retnla*, *Chil3, Ocstamp* and *Socs2* (Fig. 5A). Further, it appears that there are major changes in the gene expression profiles of WT and IL-33k.o. BMDCs already at steady-state but even more pronounced after IL-4 stimulation (Fig. 5A, B, D). Direct comparison between WT and IL-33k.o. BMDCs stimulated with IL-4 revealed that especially chemokine genes (*Ccl2*, *Ccl7*, *Ccl22* and *Ccl17*) but also *Tnfsf15* (encoding TNF-like ligand 1A, TL1A) are expressed at higher levels in WT compared to IL-33k.o. BMDCs upon IL-4 stimulation (Fig. 5A-D). Of note, CCR2 (receptor for CCL2 and CCL7) was proposed to be important for lung ILC2 expansion (*30*) and TL1A have been shown to activate ILC2s (*31*). Gene set enrichment analysis validated that IL-4-induced gene expression profiles seen in BMDCs were also detected in *ex vivo* isolated lymph node DCs that had experienced *in vivo* stimulation with IL-4, IL-13 or IL-33 (*32*) (Fig. 5E). Moreover, gene sets involved in chemokine activity, NF-κB signaling and oxidative phosphorylation were upregulated by IL-4 stimulation in WT but not in IL-33k.o. BMDCs. In contrast, gene sets related to Notch signaling and inositol phosphate metabolism were downregulated by IL-4 stimulation in WT but not in IL-33k.o. BMDCs (Fig. 5E).

**Figure 5:**
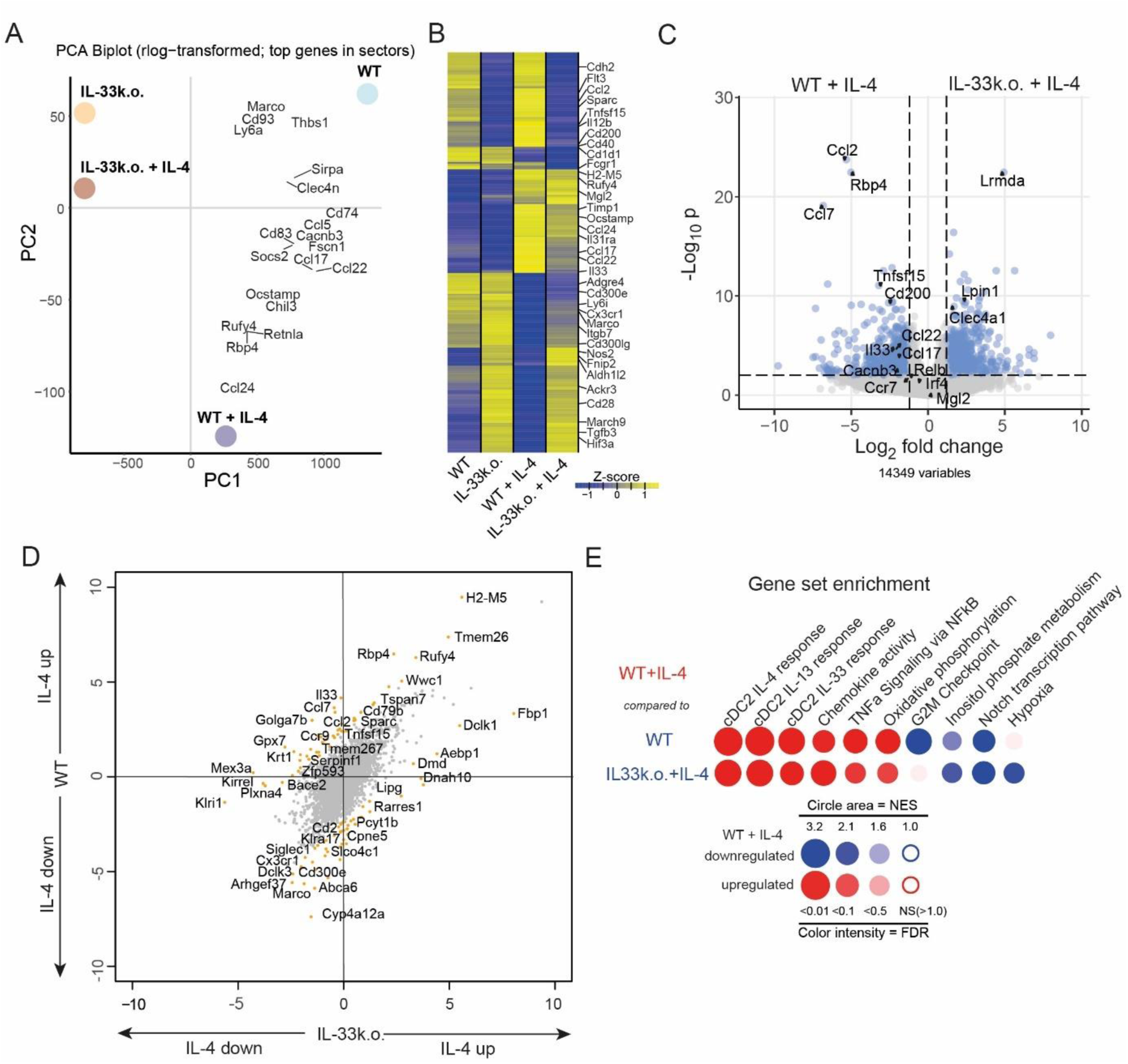
Dendritic-cell intrinsic IL-33 is important to drive the characteristic IL-4 induced signature. WT or IL-33k.o. BMDCs were stimulated with IL-4, RNA was isolated and used for bulk RNA sequencing. (A) Principal component analysis (PCA) was performed and the mean values per group of the two strongest principal components were plotted against each other and overlayed with loadings of selected genes (PCA biplot) to highlight what drives the separation (indicated by gene distance and direction relative to the crossing point of the grey lines). (B) Heatmap containing top 50 up- and top 50 downregulated genes (padj < 0.05, baseMean > 50 and log2 fold-change > 1 or < −1, respectively) for each possible comparison pair of the indicated groups. Each column is based on the mean expression values for each group. (C) Enhanced volcano plot that contrasts gene expression of WT + IL-4 and IL33k.o. + IL-4 groups. p-cutoff: 10^−3^, log2 fold-change cutoff: 1.2. (D) Log_2_-fold gene changes of the contrast WT vs WT + IL-4 on the Y-axis plotted against changes of the contrast IL-33k.o. vs IL-33k.o. + IL-4 on the X-axis. Orange spots highlight genes differentially regulated between WT and IL-33k.o. groups in an IL-4 dependent manner. E) Gene set enrichment analysis comparing transcriptomes of IL-4 stimulated WT BMDCs (WT + IL-4) with WT or IL33k.o. + IL-4 BMDCs. FDR = false discovery rate, NES = normalized enrichment score.

To further examine in which DC subset the IL-33- and IL-4-regulated transcriptional changes in BMDCs can be observed *in vivo*, we performed single-cell RNA sequencing (scRNAseq) of whole lung tissue (Fig. 6A). We subclustered the DC cluster (cluster 13), and retrieved five different populations of DCs (Fig. 6B). The comparison of DC populations before (d0) and two weeks after initial *Af* treatment (d14) revealed that *Mgl2*^+^ cDC2s as well as *Ccr7*^+^ cDC2s increase after *Af* challenge (Fig. 6C). The *Ccr7*^+^ cDC2s show high expression levels of *Ccr7*, *Cd200*, *Cacnb3*, which discriminates them from *Mgl2*^+^ cDC2s (Fig. 6C, D) and this population has already been described in other publications (*19, 33*). Expression of *Ccr7* and *Cacnb3* were reported to be important for the migratory capacity of cDC2s (*34, 35*).

**Figure 6:**
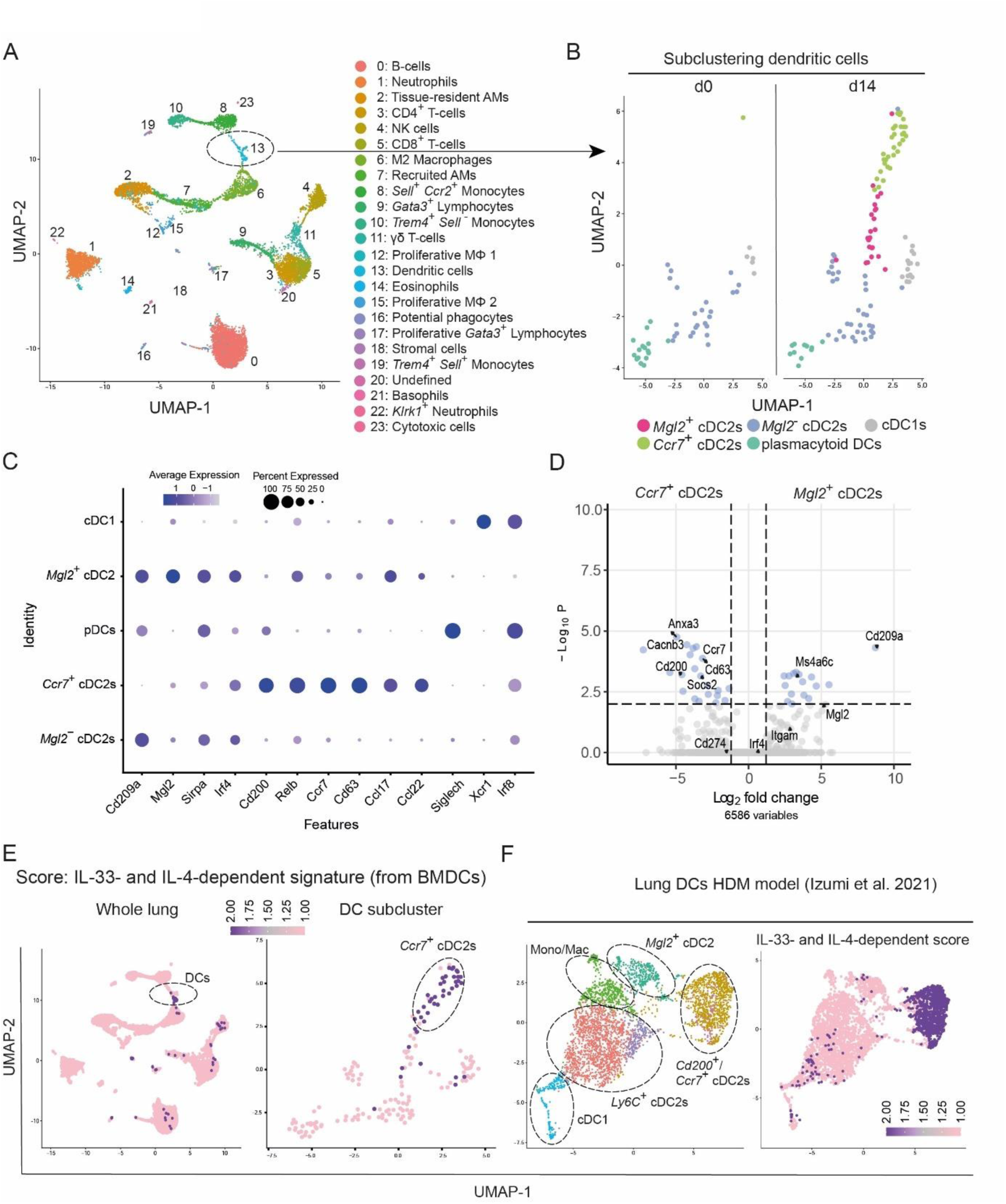
Single-cell RNA sequencing reveals different cDC2 subpopulations and indicates a role of IL-33 for their development. scRNAseq analysis was performed with whole lung tissue from untreated and *Af*-challenged WT mice. (A) Clustering revealed 23 clusters visualized by colors in a Uniform Manifold Approximation and Projection (UMAP) plot. Annotation was manually performed based on highly variable and characteristic genes. AM = alveolar macrophages. (B) Subclustering of DCs (cluster 13). Five independent DC clusters were identified. (C) Expression of selected genes in DC subclusters. (D) Volcano plot to compare differentially expressed genes between *Ccr7*^+^ cDC2s and *Mgl2*^+^ cDC2s. p-cutoff: 10^−3^, Fc cutoff: 1. (E) Gene signature score for genes exclusively upregulated in WT+IL-4 BMDCs compared to IL-33k.o.+IL-4 BMDCs (IL-33- and IL-4-dependent gene signature derived from bulk RNAseq data, p≤10^−3^, FC ≥ 1.2, Fig. 5C) was calculated for each single cell and indicated by color on top of UMAP. (F) Cluster detection and UMAP generation was performed for a dataset from house dust mite treated lungs (Izumi, Nakano, Nakano, Whitehead, Grimm, Fessler, Karmaus and Cook (*33*): Clusters and the IL-33- and IL-4-dependent gene signature score are visualized.

In order to determine whether differences of the transcriptomes observed in BMDCs are also present in DCs of the lung, we calculated a module score for the expression of significantly upregulated genes in WT+IL-4 compared to IL-33k.o.+IL-4 BMDCs. The score is separately indicated by color code for the scRNAseq data of total lung cells and the DC subcluster (Fig. 6E). A high score was mainly found in cells of the DC cluster (cluster 13) and these were mainly *Ccr7*^+^ cDC2s. To determine whether the same expression signature can also be observed in other types of allergic airway inflammation, we used a published scRNAseq dataset (*33*) in which CD11b^+^ DC subsets from lungs of house dust mite (HDM)-challenged mice were analyzed. This study could also discriminate between *Mgl2*^+^ cDC2s and *Ccr7^+^* cDC2s. When applying the BMDC-based score of IL-4- and IL-33-regulated genes on the UMAP of this data set, we observed an almost exclusive overlay with the *Ccr7*^+^ cDC2 population (Fig. 6F).

After confirming that IL-33-dependent and IL-4-induced transcriptional changes in BMDCs occur in lung cDC2s during allergic lung inflammation, we investigated whether ILC2s might respond to factors released from WT but not IL-33k.o. BMDCs upon IL-4 stimulation. Based on the bulk RNAseq data we focused on the chemokines CCL2 and CCL7 which both can be sensed by the receptor CCR2. We found *Ccr2* to be highly expressed on mRNA level in lung ILC2s in line with a previous report (*30*) (Fig. 7A, B). We confirmed CCR2 protein expression on ILC2s by flow cytometric staining (Fig. 7C). Interestingly, the mean fluorescence intensity of CCR2 staining on ILC2s decreased when WT mice had been challenged with *Af* conidia probably due to CCL2/CCL7-induced activation and internalization of CCR2. Importantly, this effect was not observed in CD11c^Cre^Irf4^fl/fl^ mice (Fig. 7C, D). Furthermore, CCR2^lo^ and CCR2^hi^ ILC2s were analyzed for the percentage of IL-13^+^ cells. This revealed that CCR2^lo^ ILC2s contained a significantly higher frequency of IL-13^+^ cells as compared to CCR2^hi^ ILC2s (Fig. 7E). These results suggest, that cDC2s mediate ligand-induced downregulation of CCR2 which correlates with increased IL-13 expression in ILC2s.

**Figure 7:**
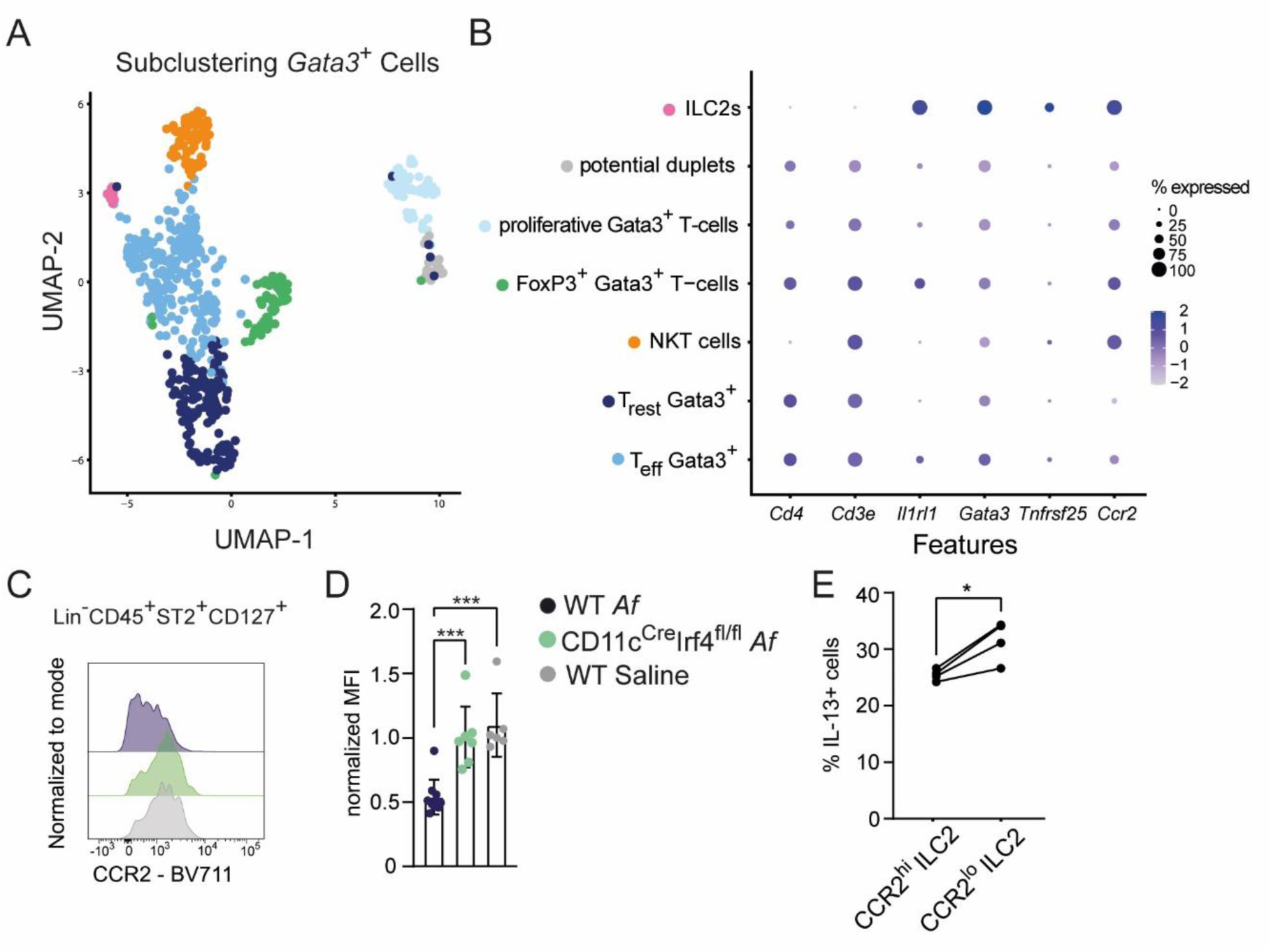
CCR2 expression is reduced on ILC2s in an *Af*- and cDC2-dependent manner. (A) Subclustering of Gata3^+^ Lymphocytes (Cluster 9+17 Fig. 6A) from the scRNAseq dataset of *Af*-treated lungs presented on a UMAP. Seven independent clusters were identified based on their expression profile. (B) Dot plot of signature genes differentiating T cells and ILC2s for the Gata3^+^ subpopulations. (C) Histograms show representative CCR2 staining on ILC2s (gated to Lin^−^CD45^+^ST2^+^CD127^+^ cells) from WT mice treated with *Af* or saline as control or CD11c^Cre^Irf4^fl/fl^ mice treated with *Af*. (D) Mean fluorescence intensity was normalized to the mean of WT mice treated with saline Bar graph shows mean±SEM of n=3-9 mice pooled from two independent experiments. One-way ANOVA was performed for statistical analysis (** p<0.01, ***p<0.001). (E) ILC2s of WT mice treated with *Af* were gated to CCR2^lo^ or CCR2^high^ and the percentage of IL-13^+^ cells was determined. Graph diplays individual values of 4 mice in two independent experiments. Paired Student’s t-test was performed for statistical analysis (* p<0.05).

Overall, these results indicate that IL-4/IL-13 stimulation of cDC2s initiates an IL-33-dependent program, including factors which can activate ILC2s, leading to ILC2 expansion in male mice which is required for lung eosinophilia during *Af*-elicited allergic lung inflammation (Fig. S4).

## Discussion

We report that ILC2s are required for eosinophilia in *Af*-induced allergic airway inflammation independently of their expression of IL-5 and in addition to Th2 cells. The mechanisms by which ILC2s accumulate in the lung and induce eosinophilia showed sex-dependent differences. Here, we propose a new model where Th2-induced, IL-33-dependent transcriptional changes in cDC2s promote ILC2 induction and eosinophilia in male mice. This leads to a similar inflammatory response in female and male mice upon *Af* challenge of the lung.

The contribution of ILC2s in *Af*-induced asthma has not been established so far. In *Alternaria alternata*- or papain-induced asthma models IL-33 release by epithelial cells is essential for ST2-dependent ILC2 activation (*36, 37*). It has been shown that *Af*-derived proteases could cleave IL-33 to induce a more potent form of IL-33 (*7*). However, no significant levels of IL-33 or TSLP were detected in *Af*-induced murine asthma (*19*). In our study we found increased ILC2s numbers in the lung even without proteolytic activity of the fungus and in the absence of the IL-33 receptor on ILC2s. Together, our results suggest that there is a mechanism independent of extracellular IL-33 which promotes accumulation of ILC2s in lung tissue. We rather showed that the ILC2 response in male mice is dependent on Th2-mediated induction of *Il33* expression in DCs. Various publications have already suggested that cDC2s are able to produce IL-33, which was important for induction of T cell responses in skin, small intestine and white adipose tissue (*28, 38, 39*). However, so far, no function for IL-33 in lung cDC2s has been described. We provide strong evidence that IL-33-dependent transcriptional changes in cDC2s are responsible for ILC2 activation in male mice. In human, it has been proposed that monocyte-derived DCs from allergic rhinitis patients produce IL-33 and that these DCs could stimulate ILC2s *in vitro* (*40*), which indicates that the mechanism we found here might also be relevant in human asthma.

The most striking IL-33-dependent transcriptional changes in BMDCs included upregulation of *Ccl2*, *Ccl7* and *Tnfsf15*. CCR2, the receptor for CCL2 and CCL7, was found to be expressed by lung ILC2s which distinguishes them from ILC2s in the small intestine. Moreover, CCR2-deficient ILC2s were less abundant in lung tissue following systemic stimulation with alarmins as compared to ILC2s from WT mice (*30*). Our results indicate that CCR2 on lung ILC2s is downregulated in an *Af*- and cDC2-dependent manner, hinting towards activation and internalization of CCR2. Beside CCR2, the engaging receptor of TNFSF15/TL1A, DR3 has been shown to be important for ILC2 function, survival, and expansion in a papain challenge model (*31*). Furthermore, DR3 was suggested as costimulatory receptor in ILC2s in patients suffering from eosinophilic asthma (*41*). As the mechanism of ILC2 induction by IL-33^+^ cDC2s is primarily important in male mice, cDC2-derived molecules might counteract the inhibitory androgen receptor (AR) signaling in ILC2s which is believed to be the main reason for suppression of the ILC2 response in male mice compared to females. Interestingly, in prostate cancer CCR2 signaling was shown to specifically counteract AR signaling (*42*). Whether there is a similar mechanism in lung ILC2 should be further investigated. Taken together, we found that cDC2s can produce stimulatory molecules for ILC2s in an IL-4-induced and IL-33-dependent manner.

Additionally, we found the IL-4-plus IL-33-regulated gene signature to be expressed especially in a *Ccr7*^+^ cDC2 population. Of note, we found *Il33* expression already in MGL2^+^ cDC2s, however, the subsequently IL-33-regulated genes appeared mostly in *Ccr7*^+^ cDC2s which is in line with a publication by Izumi et al. (*33*) showing that *Ccr7*^+^ cDC2s originate from *Mgl2*^+^ cDC2s. Other reports provided evidence that IL-4/IL-13 signaling via STAT6 can induce migratory DCs (*12, 43, 44*). Together, this indicates that transition from *Mgl2*^+^ cDC2s to *Ccr7*^+^ migratory cDC2s is partly dependent on IL-4-induced and IL-33-dependent transcriptional changes.

ILC2s have been shown to play an important non-redundant role for eosinophilia in allergic airway inflammation, especially in models with high IL-33 levels in the lung (*14*). Despite the fact that ILC2s only constitute a minor population of cytokine-producing cells in the *Af*-challenged lung (*19*), we found that they play an important role for eosinophilia in addition to Th2 cells. Although ILC2-derived IL-5 is considered to be important for eosinophil homeostasis as well as eosinophilia in inflammation (*9*), mice with ILC2-specific deletion of IL-5 have not been investigated in this regard. To this end, we used a chimeric mouse approach and surprisingly IL-5 deficiency in ILC2s had no effect on eosinophilia in the *Af*-induced lung inflammation. Possibly, IL-5 is still produced in decent amounts from other sources, including T cells, which seems to be sufficient for eosinophilia in this context. We rather found male ILC2s as essential source of IL-4/IL-13 which promoted CCL24 production in macrophages and this led to efficient recruitment of eosinophils into the lung. This result also indicates that such sex-specific differences in eotaxin production might be relevant for eosinophilia in other settings of allergic asthma. As male and female mice mount a similar lung eosinophilia upon *Af* challenge, it remains to be investigated how female mice compensate for the lower CCL24 expression.

In conclusion, our study provides strong evidence for a novel Th2-IL33^+^cDC2-ILC2-eosinophil axis which is required in male mice for *Af*-induced allergic lung inflammation. These findings contribute to the understanding of sex-dependent differences in asthma pathology and indicate that new approaches for future development of sex-dependent treatments of asthma patients should be considered.

## Material and Methods

### Mice

Tie2^Cre^RORα^fl/fl^ (ILC2k.o.)(*45, 46*), IL-33-deficient (IL-33k.o., originally from the trans-NIH Knock-Out Mouse Project (KOMP) Repository, www.komp.org), IL-5-deficient (Red5)(*9*), and ST2-deficient (Il1rl1k.o.)(*47*) mice were kindly provided by Stefan Wirtz. KLRG1^Cre^4-13^fl/fl^ mice were generated by crossing KLRG1^Cre^ mice (*48*) to Rosa26^LSL-tdtomato^ (*49*) and conditional IL-4/IL-13-deficient (4-13^fl/fl^)(*50*) mice. Clec9a^Cre^Irf4^fl/fl^ (cDC2k.o.)(*51*) mice were generated and kindly provided by Barbara Schraml. CD11c^Cre^Irf4^fl/fl^ (*52, 53*), CD4^Cre^4-13^fl/fl^ (*54*) and STAT6-deficient (*55*) mice have been described previously. All mice were used on C57BL/6 background, housed under SPF conditions and used for experiments at an age of 7-12 weeks at the start of the experiments which were performed with approval by the Government of Lower Franconia according to the German animal protection law and European Union guidelines (Directive 2010/63/EU).

### Generation of mixed bone marrow chimeras

Bone marrow from donor mice was mixed in either a 1:4 (Red5 or WT:ILC2k.o.; IL-33k.o. or WT:cDC2k.o.) or 1:1 (ST2k.o.-CD45.2:WT-CD45.1) ratio. Recipient mice were irradiated with 11 Gy and then received 2×10^6^ cells of the respective bone marrow mixture. The mice were treated for four weeks with antibiotics-containing drinking water and were used after eight weeks for experiments.

### Intranasal treatment with *Af* conidia

Mice were intranasally treated with freshly isolated *Aspergillus fumigatus* ATCC 46645 or *Aspergillus fumigatus* prtTΔ (on ATCC 46645 background) (*56*) conidia as described previously(*5*). The mice were analyzed on day 14 of experiments.

### Organ sampling, preparation of single-cell suspension and intracellular staining

Preparation of single-cell suspension from the lung was performed as described previously(*5*). For flow cytometric staining, single cell suspensions were first incubated with rat anti-mouse CD16/CD32 antibody (clone 2.4G2, BioXcell) to prevent unspecific staining. The following antibodies were applied for 20 min at 4°C: Anti-CD3-PE-Cy7 (17A2), anti-CD4-PerCP-Cy5.5 (RM4-5), anti-CD8-PE-Cy7 (53-6.7), anti-CD11c PE-Cy7/ BV786 (N4/18), anti-CD45 APC-Cy7 (30-F11), anti-CD64 FITC/ PE-Cy7 (X54-5/7.1), anti-CD127 PerCP-Cy5.5 (A7R34), anti-CD172a PerCP-Cy5.5 (P84), anti-CD301b/Mgl2 AF647/ PE-Dazzle (URA1), anti-KLRG1 PerCP-Cy5.5 (2F1) all from Biolegend (San Diego, CA). Anti-CD4 PE (RM4-5), anti-CD11b PE-Cy7 (M1/70), anti-CD45R PE-Cy7 (RA3-6B2), anti-CD101 PE (Moushi101), anti-CD127 BV786 (SB/199), anti-FcεRI alpha PE-Cy7 (Mar-1), anti-MerTK PE-Cy7 (DS5MMER) all from eBioscience/ ThermoFisher Scientific (Waltham, MA). Anti-NK1.1 PE-Vio770 (PK136), anti-NKp46 PE-Vio770 (REA815) from Miltenyi Biotec (Bergisch Gladbach, Germany). Anti-CD4 BUV737 (RM 4-5), anti-CD11c BUV737 (N4/18), anti-CD45.1 BUV737 (A20), anti-CD45.2 BV786 (104), anti-CD192 BV711 (475301) anti-KLRG1 BV711 (2F1), anti-MHC-II BUV395 (2G9), anti-Siglec-F BV421 (E50-2440), anti-ST2 BV421 (U29-93), all from BD Biosciences (San Jose, CA).

For intracellular staining of cytokines, cells were restimulated with PMA (50 ng/ml) and ionomycin (2 µg/ml) for 1.5 h followed by incubation with BD GolgiPlug (1:1000) for 2 h. The cells were first stained with surface antibodies, subsequently the BD Cytofix/Cytoperm Fixation/Permeabilization Kit was used according to the manufacturer’s protocol. Following intracellular antibodies were stained over night at 4°C: anti-IL-4-PE (11B11), anti-IL-5-APC (TRFK5) both from BioLegend. Anti-IL-13-FITC or PE (eBio13A) from eBioscience/Thermo Fisher Scientific).

For intracellular staining of transcription factors, after initial staining of surface antibodies, the Foxp3/Transcription Factor Staining Buffer Set from eBioscience/Thermo Fisher Scientific was used following the manufacturer’s protocol for fixation of cells. The following antibodies were incubated with the cells over night at 4°C: anti-Gata3 BUV395 (L50-823, BD Biosciences), anti-FoxP3 APC (FJK-16s, eBioscience/Thermo Fisher Scientific.

Flow cytometric measurements were performed with a BD LSRFortessa instrument and analyses were performed with FlowJo software (Version X, Treestar, Ashland, OR).

### Cell Sorting

For sorting of Mgl2^+^ cDC2s, single-cell suspensions were incubated with anti-Siglec-F-biotin (E50-2440, BD Bioscience) and anti-CD64-biotin (X54-5/7.1, BioLegend) and negative selection with Anti-Biotin MicroBeads (Miltenyi Biotec) was performed according to the manufacturer’s protocol. Further, the negative selected cells or total lung cells (for sorting of macrophages) were stained with the indicated fluorophore-labelled antibodies and sorting was performed using a S3 Cell Sorter (Bio-Rad, Hercules, CA).

### BM-derived DC culture

BM of hind leg bones of WT, IL-33k.o., or STAT6k.o. mice was isolated. Following red blood cell lysis using ACK lysis buffer, cells were cultured at a concentration of 2×10^5^ cells/ml for 4 days and further 6 days at a concentration of 1×10^6^ cells/ml in RPMI1640 medium supplemented with 5% GM-CSF-containing supernatant of X63 cells. At day 10 of culture, cells were harvested and 5×10^5^ cells were seeded in 200 µl RPMI1640, optionally containing IL-4 (R&D Systems, 5 ng/ml), in a 96-well plate for 24 h. RNA of cultured BMDCs was isolated using the RNeasy Micro Kit (Qiagen, Hilden, Germany) following the provided protocol. Bulk RNA sequencing was performed by Novogen (Munich, Germany).

### Quantitative real-time PCR

RNA from whole lung tissue or sorted cells was isolated with TRIzol reagent (Thermo Fisher Scientific) and chloroform extraction. cDNA was generated using the SuperScript III Reverse Transcriptase (Thermo Fisher Scientific) and oligo(dT)primers. Real-time PCR was performed using SYBR Select Master Mix (Thermo Fisher Scientific) and the following primers: Hprt-fwd 5’-GTT GGA TAC AGG CCA GAC TTT GTT-3’; Hprt-rev 5’-GAG GGT AGG CTG GCC TAT AGG CT-3’; Il4-fwd 5’-ACT TGA GAG AGA TCA TCG GCA-3’; Il4-rev (5’-AGC TCC ATG AGA ACA CTA GAG TT -3’; Il5-fwd 5’- TCT GTA CTC ATC ACA CCA AG -3’; Il5-rev 5’- AGG ATG CTT CTG CAC TTG AG -3’; Il13-fwd 5’- TGC TTT GTG TAG CTG AGC AG -3‘; Il13-rev 5‘- GCA GTC CTG GCT CTT GCT TG -3’; Il33-fwd 5’-TCC AAC TCC AAG ATT TCC CCG – 3’; Il33-rev 5’- CAT GCA GTA GAC ATG GCA GAA -3’; Ccl11-fwd 5’- TTC CAT CTG TCT CCC TCC ACC AT -3’; Ccl11-rev 5’- CCT GGT CTT GAA GAC TAT GGC TTT CA -3’; Ccl24-fwd 5’- TCT TCC CCA TAG ATT CTG TGA CCA -3’; Ccl24-rev 5’- GTT TTT GTA TGT GCC TCT GAA CCC -3’. Measurements were performed using a ViiA 7 Real-Time PCR System (Applied Biosystems/Thermo Fisher Scientific). C_t_ values of targets were normalized to C_t_ values of *Hprt*.

### Bulk RNAseq analysis

RNA was sent to Novogene Europe (Munich) for sequencing as a service. There, library preparation was performed with NEBNext® UltraTM RNA Library Prep Kit for Illumina (NEB, Frankfurt, Germany) following manufacturer’s recommendations and paired-end sequencing was performed on an Illumina NovaSeq6000. Filtered clean reads were aligned to reference genome mm10 using STAR (v2.6.1d)(*57*). FeatureCounts (v1.5.0-p3) was used to generate counts used for downstream analysis(*58*). Data were loaded into R (3.5.3; The R Foundation for Statistical Computing, Vienna, Austria) and genes with zero counts were removed. Deseq2 (1.20.0)(*59*) was used to normalize counts and perform differential expression analysis without further shrinkage. Heatmaps of log2 normalized counts were drawn with the gplots package. For gene set enrichment Bubble GUM software (1.3.19)(*60*) based on GSEA(*61*) with normalized count values as input and recommended settings (default values but permutation type set to “gene_set”) was used. Differentially expressed genes were visualized with the “Enhanced Volcano” package (https://github.com/kevinblighe/EnhancedVolcano). For the GSEA analysis selected gene sets of MSigDB database version 2025.1 for *Mus musculus* were used. Data is available via ArrayExpress accession E-MTAB-15732.

### Single-cell RNA sequencing

Two C57BL/6 mice per time point (d0, d14) of *Aspergillus fumigatus* treatment were analysed for single-cell sequencing. Single-cell suspensions from lungs were processed with the dead cell removal kit (Miltenyi Biotec), labelled with the BD Ms Single Cell Sample Multiplexing Kit (BD Bioscience) and stained for viable cells with DRAQ7 and Calcein AM (both Thermo Fisher Scientific). Then cDNA of single cells was captured using the BD Rhapsody Single-Cell Analysis System following the provided protocol. Sample Tag and Whole Transcriptome Analysis libraries were prepared according to the manufacturer’s instructions. In the process, quantity was measured using the Qubit Fluorometer (Thermo Fisher Scientific) and quality control was performed using the Agilent 4200 TapeStation system with the Agilent High Sensitivity D5000 ScreenTape Assay (Agilent). Sequencing was performed at a NovaSeq 6000 instrument (Illumina, San Diego, CA). Data was processed through BD Rhapsody WTA Analysis Pipeline version 1.12 and loaded into R for further analysis. To be included in the analysis cells needed to have >2500 UMIs, >1000 genes detected per cell, <10% mitochondrial reads, and a novelty >0.8 (log10 of gene number divided by log10 of UMIs). Data normalization, differential expression analysis, clustering, and dimensional reduction (UMAP based on top 40 principal components) were performed in Seurat (version 5.0.1)(*62*) under R (version 4.2.0). Gene set scores were calculated in Seurat with the corresponding function(*63*). Data is available via ArrayExpress accession E-MTAB-16136.

### Statistical analysis

Graph Pad Prism 10 (GraphPad Software, Boston, MA) was used for statistical analysis. Statistical tests used are mentioned in figure legends. Outlier removal was legitimated based on ROUT test. Bar graphs show the mean±SEM from samples of at least two independent experiments.

## Acknowledgments

We thank the sequencing core unit at the Department of Human Genetics of the FAU, especially Arif Ekici. We thank Daniela Döhler and Michaela Dümig for technical support, and members of the Voehringer lab for helpful discussions.

## Author contributions

L.-M.G., A.R., D.V. designed experiments, L.-M.G., D.R., A.R., and K.C. performed experiments, L.-M.G. and D.R. performed computational analysis, S.K. provided critical reagents and organisms, S.W. and B.U.S. provided mouse strains and bone marrow, L.-M.G., D.R. and D.V. wrote the manuscript.

## Data availability

RNA-Sequencing data was uploaded to ArrayExpress. Bulk RNA Sequencing data is available via ArrayExpress accession E-MTAB-15732. Single-cell RNA Sequencing data is available via ArrayExpress accession E-MTAB-16136.

## References

1. G. B. D. Asthma, C. Allergic Diseases, Global, regional, and national burden of asthma and atopic dermatitis, 1990-2021, and projections to 2050: a systematic analysis of the Global Burden of Disease Study 2021. Lancet Respir Med 13, 425–446 (2025).

2. J. P. Latge, G. Chamilos, Aspergillus fumigatus and Aspergillosis in 2019. Clin Microbiol Rev 33, (2019).

3. R. Agarwal, V. Muthu, I. S. Sehgal, Relationship between Aspergillus and asthma. Allergol Int 72, 507–520 (2023).

4. J. Furlong-Silva, P. C. Cook, Fungal-mediated lung allergic airway disease: The critical role of macrophages and dendritic cells. PLoS Pathog 18, e1010608 (2022).

5. A. Dietschmann, S. Schruefer, S. Krappmann, D. Voehringer, Th2 cells promote eosinophil-independent pathology in a murine model of allergic bronchopulmonary aspergillosis. Eur J Immunol 50, 1044–1056 (2020).

6. S. G. Smith et al., Increased numbers of activated group 2 innate lymphoid cells in the airways of patients with severe asthma and persistent airway eosinophilia. J Allergy Clin Immunol 137, 75–86 e78 (2016).

7. C. Cayrol et al., Environmental allergens induce allergic inflammation through proteolytic maturation of IL-33. Nat Immunol 19, 375–385 (2018).

8. I. Martinez-Gonzalez, C. A. Steer, F. Takei, Lung ILC2s link innate and adaptive responses in allergic inflammation. Trends Immunol 36, 189–195 (2015).

9. J. C. Nussbaum et al., Type 2 innate lymphoid cells control eosinophil homeostasis. Nature 502, 245–248 (2013).

10. K. F. Troch et al., Group 2 innate lymphoid cells are a non-redundant source of interleukin-5 required for development and function of murine B1 cells. Nat Commun 15, 10566 (2024).

11. J. Wan et al., HMGB1-induced ILC2s activate dendritic cells by producing IL-9 in asthmatic mouse model. Cell Immunol 352, 104085 (2020).

12. T. Y. Halim et al., Group 2 innate lymphoid cells are critical for the initiation of adaptive T helper 2 cell-mediated allergic lung inflammation. Immunity 40, 425–435 (2014).

13. R. K. Gurram et al., Crosstalk between ILC2s and Th2 cells varies among mouse models. Cell Rep 42, 112073 (2023).

14. K. J. Jarick et al., Non-redundant functions of group 2 innate lymphoid cells. Nature 611, 794–800 (2022).

15. Y. Yasuda, T. Nagano, K. Kobayashi, Y. Nishimura, Group 2 Innate Lymphoid Cells and the House Dust Mite-Induced Asthma Mouse Model. Cells 9, (2020).

16. S. Laffont et al., Androgen signaling negatively controls group 2 innate lymphoid cells. J Exp Med 214, 1581–1592 (2017).

17. E. Blanquart et al., Targeting androgen signaling in ILC2s protects from IL-33-driven lung inflammation, independently of KLRG1. J Allergy Clin Immunol 149, 237–251 e212 (2022).

18. R. Karkout et al., Female-specific enhancement of eosinophil recruitment and activation in a type 2 innate inflammation model in the lung. Clin Exp Immunol 216, 13–24 (2024).

19. P. C. Cook et al., Mgl2(+) cDC2s coordinate fungal allergic airway type 2, but not type 17, inflammation in mice. Nat Commun 16, 928 (2025).

20. M. Kopf et al., IL-5-deficient mice have a developmental defect in CD5+ B-1 cells and lack eosinophilia but have normal antibody and cytotoxic T cell responses. Immunity 4, 15–24 (1996).

21. S. Huber, R. Hoffmann, F. Muskens, D. Voehringer, Alternatively activated macrophages inhibit T-cell proliferation by Stat6-dependent expression of PD-L2. Blood 116, 3311–3320 (2010).

22. B. U. Schraml et al., Genetic tracing via DNGR-1 expression history defines dendritic cells as a hematopoietic lineage. Cell 154, 843–858 (2013).

23. Y. Gao et al., Control of T helper 2 responses by transcription factor IRF4-dependent dendritic cells. Immunity 39, 722–732 (2013).

24. J. W. Williams et al., Transcription factor IRF4 drives dendritic cells to promote Th2 differentiation. Nat Commun 4, 2990 (2013).

25. B. W. Li et al., T cells are necessary for ILC2 activation in house dust mite-induced allergic airway inflammation in mice. Eur J Immunol 46, 1392–1403 (2016).

26. O. Liska et al., TFLink: an integrated gateway to access transcription factor-target gene interactions for multiple species. Database (Oxford) 2022, (2022).

27. Z. Wang et al., SUMOylated IL-33 in the nucleus stabilizes the transcription factor IRF1 in hepatocellular carcinoma cells to promote immune escape. Sci Signal 16, eabq3362 (2023).

28. J. M. Inclan-Rico et al., MrgprA3 neurons drive cutaneous immunity against helminths through selective control of myeloid-derived IL-33. Nat Immunol 25, 2068–2084 (2024).

29. J. Helft et al., GM-CSF Mouse Bone Marrow Cultures Comprise a Heterogeneous Population of CD11c(+)MHCII(+) Macrophages and Dendritic Cells. Immunity 42, 1197–1211 (2015).

30. M. Zhao et al., Maturation and specialization of group 2 innate lymphoid cells through the lung-gut axis. Nat Commun 13, 7600 (2022).

31. X. Yu et al., TNF superfamily member TL1A elicits type 2 innate lymphoid cells at mucosal barriers. Mucosal Immunol 7, 730–740 (2014).

32. A. Cui et al., Dictionary of immune responses to cytokines at single-cell resolution. Nature 625, 377–384 (2024).

33. G. Izumi et al., CD11b(+) lung dendritic cells at different stages of maturation induce Th17 or Th2 differentiation. Nat Commun 12, 5029 (2021).

34. M. S. Woo et al., Calcium channel beta3 subunit regulates ATP-dependent migration of dendritic cells. Sci Adv 9, eadh1653 (2023).

35. A. Meloun, B. Leon, Beyond CCR7: dendritic cell migration in type 2 inflammation. Front Immunol 16, 1558228 (2025).

36. P. M. Topczewska et al., ILC2 require cell-intrinsic ST2 signals to promote type 2 immune responses. Front Immunol 14, 1130933 (2023).

37. R. J. Snelgrove et al., Alternaria-derived serine protease activity drives IL-33-mediated asthma exacerbations. J Allergy Clin Immunol 134, 583–592 e586 (2014).

38. L. Y. Hung et al., Cellular context of IL-33 expression dictates impact on anti-helminth immunity. Sci Immunol 5, (2020).

39. S. Soedono et al., Obese visceral adipose dendritic cells downregulate regulatory T cell development through IL-33. Front Immunol 15, 1335651 (2024).

40. Y. Q. Peng et al., Effects of myeloid and plasmacytoid dendritic cells on ILC2s in patients with allergic rhinitis. J Allergy Clin Immunol 145, 855–867 e858 (2020).

41. K. Machida et al., The Role of the TL1A/DR3 Axis in the Activation of Group 2 Innate Lymphoid Cells in Subjects with Eosinophilic Asthma. Am J Respir Crit Care Med 202, 1105–1114 (2020).

42. Q. Tang, B. Cheng, R. Dai, R. Wang, The Role of Androgen Receptor in Cross Talk Between Stromal Cells and Prostate Cancer Epithelial Cells. Front Cell Dev Biol 9, 729498 (2021).

43. T. Y. Halim et al., Group 2 innate lymphoid cells license dendritic cells to potentiate memory TH2 cell responses. Nat Immunol 17, 57–64 (2016).

44. S. Lee et al., STAT6 inhibitory peptide reduces dendritic cell migration to the lymph nodes to control Th2 adaptive immunity in the mouse lung. Eur J Immunol 49, 157–169 (2019).

45. L. Knipfer et al., A CCL1/CCR8-dependent feed-forward mechanism drives ILC2 functions in type 2-mediated inflammation. J Exp Med 216, 2763–2777 (2019).

46. Y. Omata et al., Group 2 Innate Lymphoid Cells Attenuate Inflammatory Arthritis and Protect from Bone Destruction in Mice. Cell Rep 24, 169–180 (2018).

47. K. Hoshino et al., The absence of interleukin 1 receptor-related T1/ST2 does not affect T helper cell type 2 development and its effector function. J Exp Med 190, 1541–1548 (1999).

48. D. Herndler-Brandstetter et al., KLRG1(+) Effector CD8(+) T Cells Lose KLRG1, Differentiate into All Memory T Cell Lineages, and Convey Enhanced Protective Immunity. Immunity 48, 716–729 e718 (2018).

49. L. Madisen et al., A robust and high-throughput Cre reporting and characterization system for the whole mouse brain. Nat Neurosci 13, 133–140 (2010).

50. D. Voehringer, D. Wu, H. E. Liang, R. M. Locksley, Efficient generation of long-distance conditional alleles using recombineering and a dual selection strategy in replicate plates. BMC Biotechnol 9, 69 (2009).

51. N. Salei et al., Selective depletion of a CD64-expressing phagocyte subset mediates protection against toxic kidney injury and failure. Proc Natl Acad Sci U S A 118, (2021).

52. M. L. Caton, M. R. Smith-Raska, B. Reizis, Notch-RBP-J signaling controls the homeostasis of CD8-dendritic cells in the spleen. J Exp Med 204, 1653–1664 (2007).

53. U. Klein et al., Transcription factor IRF4 controls plasma cell differentiation and class-switch recombination. Nat Immunol 7, 773–782 (2006).

54. K. Oeser, C. Schwartz, D. Voehringer, Conditional IL-4/IL-13-deficient mice reveal a critical role of innate immune cells for protective immunity against gastrointestinal helminths. Mucosal Immunol 8, 672–682 (2015).

55. M. H. Kaplan, U. Schindler, S. T. Smiley, M. J. Grusby, Stat6 is required for mediating responses to IL-4 and for development of Th2 cells. Immunity 4, 313–319 (1996).

56. A. Bergmann, T. Hartmann, T. Cairns, E. M. Bignell, S. Krappmann, A regulator of Aspergillus fumigatus extracellular proteolytic activity is dispensable for virulence. Infect Immun 77, 4041–4050 (2009).

57. A. Dobin et al., STAR: ultrafast universal RNA-seq aligner. Bioinformatics 29, 15–21 (2013).

58. Y. Liao, G. K. Smyth, W. Shi, featureCounts: an efficient general purpose program for assigning sequence reads to genomic features. Bioinformatics 30, 923–930 (2014).

59. M. I. Love, W. Huber, S. Anders, Moderated estimation of fold change and dispersion for RNA-seq data with DESeq2. Genome Biol 15, 550 (2014).

60. L. Spinelli, S. Carpentier, F. Montanana Sanchis, M. Dalod, T. P. Vu Manh, BubbleGUM: automatic extraction of phenotype molecular signatures and comprehensive visualization of multiple Gene Set Enrichment Analyses. BMC Genomics 16, 814 (2015).

61. A. Subramanian et al., Gene set enrichment analysis: a knowledge-based approach for interpreting genome-wide expression profiles. Proc Natl Acad Sci U S A 102, 15545–15550 (2005).

62. Y. Hao et al., Dictionary learning for integrative, multimodal and scalable single-cell analysis. Nat Biotechnol 42, 293–304 (2024).

63. I. Tirosh et al., Dissecting the multicellular ecosystem of metastatic melanoma by single-cell RNA-seq. Science 352, 189–196 (2016).

