## Supplemental Figures for "*Il33* expressing cDC2s promote expansion of ILC2s and eosinophilia in fungal airway inflammation in male mice"

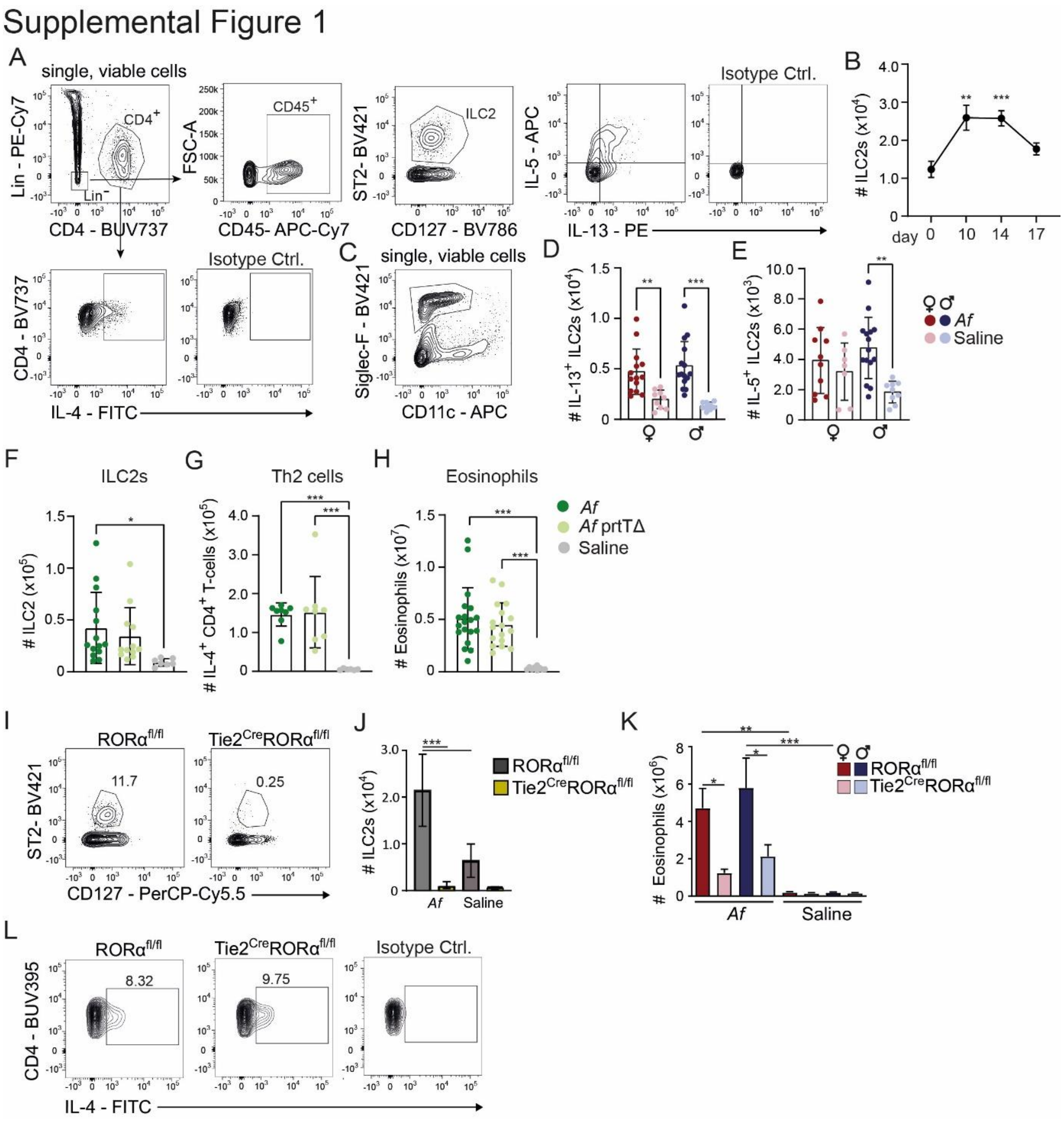


**Fig. S1: Gating strategy for flow cytometry, comparison of *Af* WT and *Af* prtTΔ and confirmation of ILC2-deficiency in Tie2 $^{Cre}$Rora$^{fl/fl}$ mice.** (A) Gating strategy for lung ILC2s (pre-gated to lymphocytes, single cells and viable cells) and IL-$4^+$ T-cells, Lin: CD3, CD8, CD11b, CD11c, CD45R, FcεRI alpha, NK1.1, NKp46. (B) Graph shows total ILC2 numbers at different time points after *Aspergillus fumigatus* treatment. (C) Gating strategy of eosinophils (pre-gated to single cells and viable cells). (D, E) Total number of IL-$13^+$ (D) and IL-$5^+$ (E) ILC2s separated by sex in the lungs of WT mice. (F-H) WT mice were treated with *Af* WT strain, an *Af* mutant strain which lacks extracellular protease activity (*Af* prtTΔ) or saline (Ctrl). Total numbers of ILC2s (F), IL-$4^+$ T-cells (G) and eosinophils (H) were analysed in the lung at d14 of treatment. (I-L) Tie2$^{Cre}$Rora$^{fl/fl}$ mice were treated with *Af* and analyzed on day 14. (I) Representative flow cytometric gating of ILC2s. (J) Quantitatively analyzed numbers of ILC2s in (I) are displayed (K) Total numbers of eosinophils in female and male mice are shown in the bar graph. (L) Gating strategy for IL-$4^+$ T-cells is displayed. (B, D, E, F-H, J, K) Graphs display mean±SEM of $n \geq 6$ (B), $n \geq 6$ (D, E), $n \geq 7$ (E), $n \geq 6$ (F-H), $n \geq 4$ (J, K) mice per group in 3 (B), 7 (D, E), 3 (F-H), 2 (J) and 4 (K) independent experiments. One-way ANOVA was performed for statistical testing (* $p<0.05$, ** $p<0.01$, *** $p<0.001$). Only relevant statistical significances are indicated.

### Supplemental Figure 2

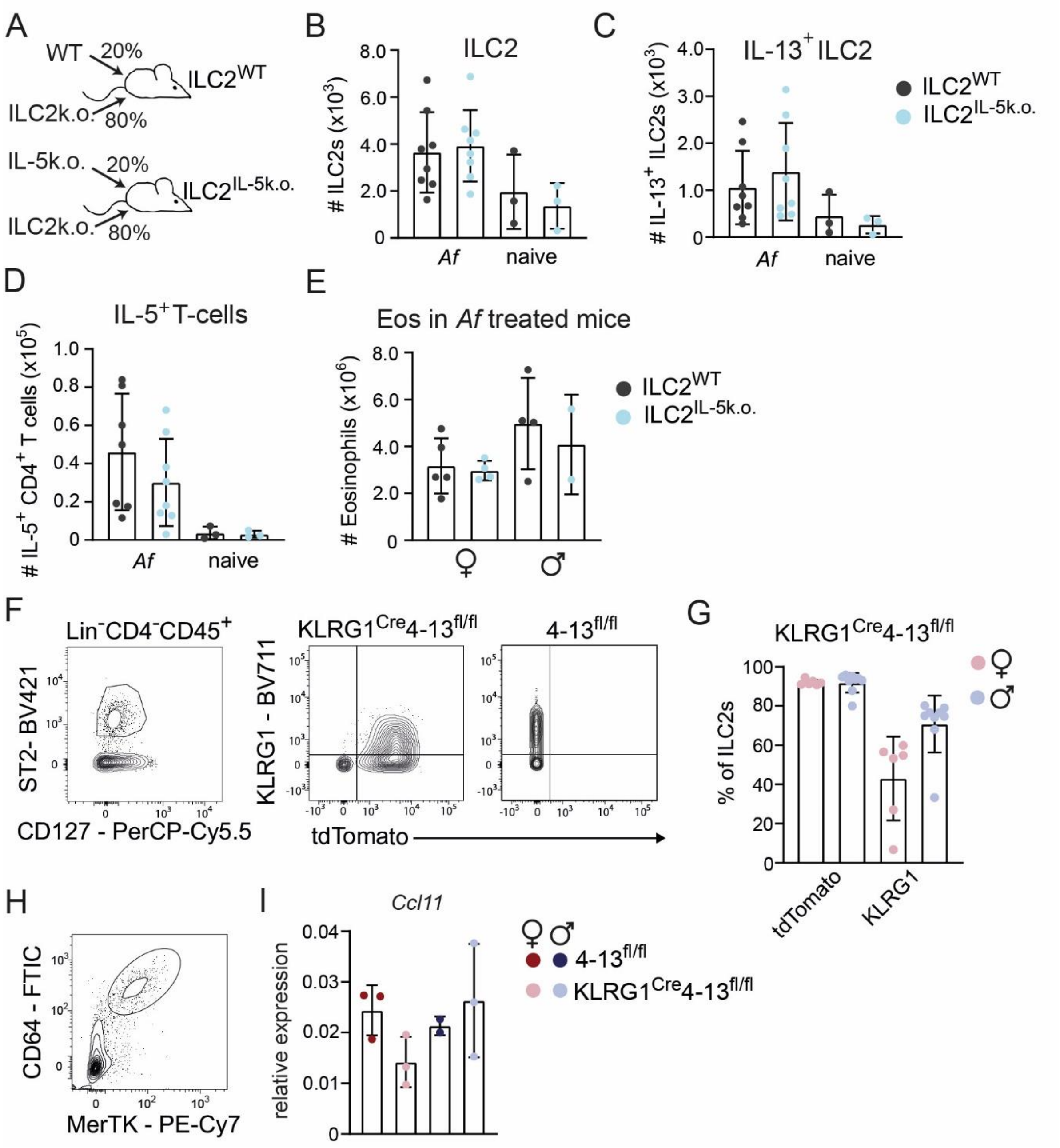


**Fig. S2: Analysis of cytokine-deficient ILC2 mouse models**: (A) Lethally irradiated ILC2-deficient mice were reconstituted with 20% WT or 20% Red5 (IL-5k.o.) and 80% ILC2k.o. (Tie2$^{Cre}$RORα$^{fl/fl}$) bone marrow. Total number of ILC2s (B), IL-13$^{+}$ ILC2s (C), IL-5$^{+}$ T-cells (D) is displayed as mean±SEM in the graphs. (E) Total number of eosinophils separated by sex in *Af*-treated mixed BM chimeras is shown in the bar graph. (F) Exemplary gating strategy of ILC2s as well as expression of KLRG1 and tdTomato is displayed in the flow cytometric plots. (G) Percentages of tdTomato$^{+}$ and KLRG1$^{+}$ ILC2s in (F) is displayed in the bar graph. (H) Flow cytometric plot indicates flow cytometric gating strategy for sorting macrophages. (I) Quantitative RT-PCR of *Ccl11* from total lung tissue normalized to *Hprt*.

#### Supplemental Figure 3

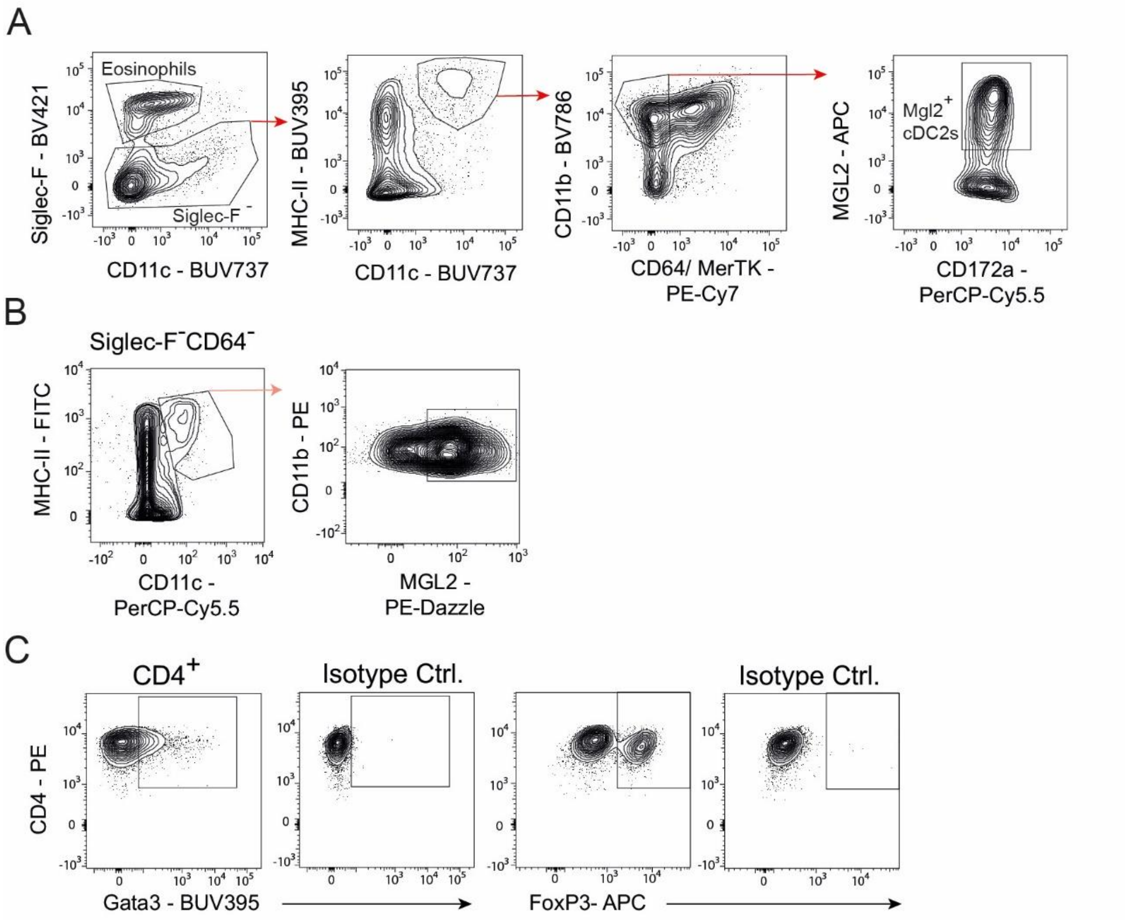


**Fig. S3: Gating strategies for flow cytometry and flow cytometry-based cell sorting.** (A) Gating strategy for eosinophils as well as Mgl2$^{+}$ cDC2s is displayed in flow cytometric plots. (B) Flow cytometric plots indicate gating for Mgl2$^{+}$ cDC2 cell sorting. Cells were pre-sorted for CD64$^{-}$, Siglec-F$^{-}$ negative cells with anti-biotin micro beads. (C) Gating of Gata3$^{+}$ and FoxP3$^{+}$ T-cells is indicated in flow cytometric plots.

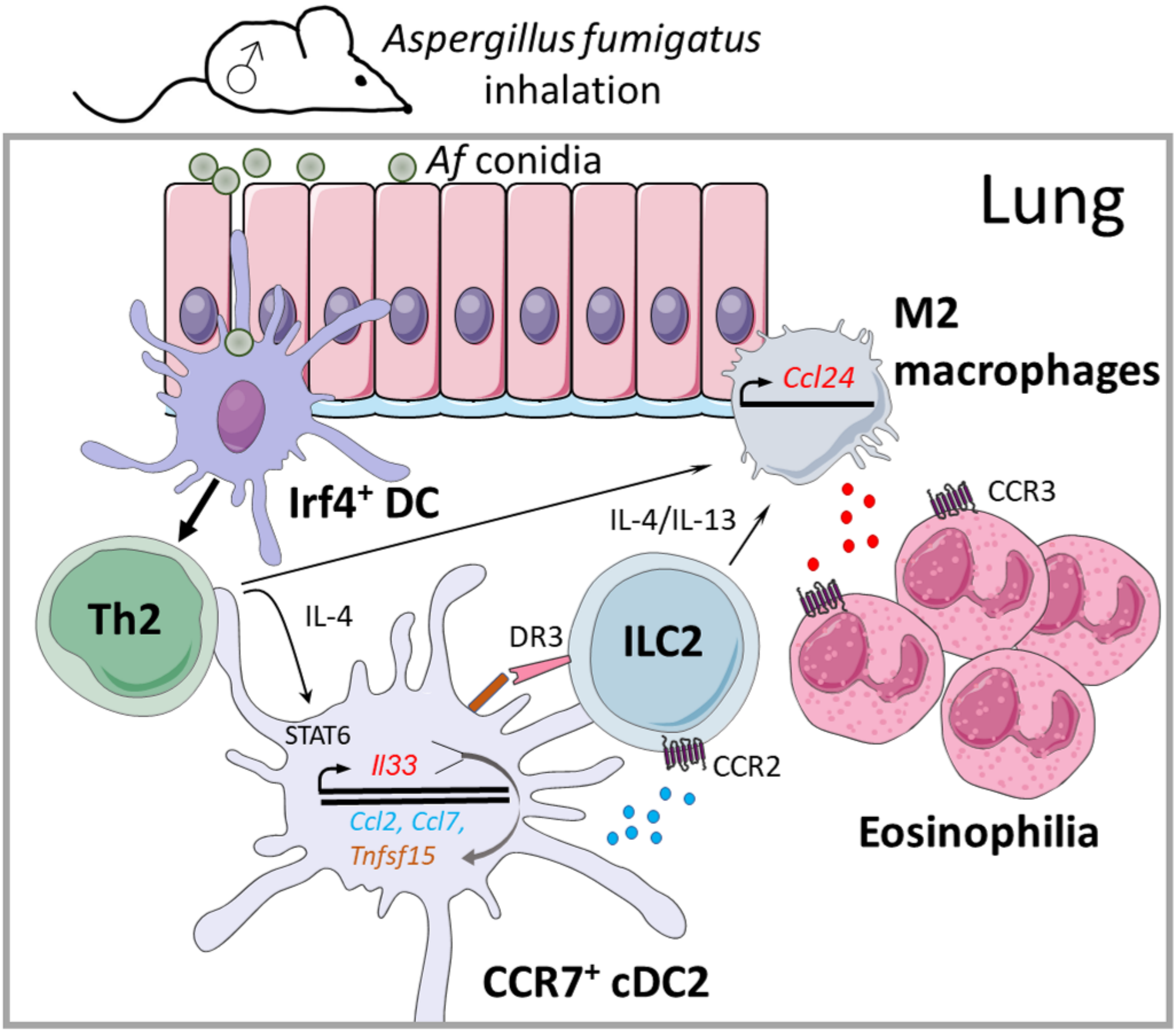


**Fig. S4: IL-4-induced and IL-33-dependent factors of cDC2s drive ILC2 expansion necessary for lung eosinophilia in male mice upon *Aspergillus fumigatus* induced allergic inflammation.** In summary, our data suggest a novel mechanism in male mice, whereby Th2 cells feedback on cDC2s and elicit an IL-4-induced and IL-33-dependent transcriptional profile. This leads to expression of ILC2 stimulatory factors which promote ILC2 expansion in male animals. ILC2-derived IL-4/IL-13 then induces Ccl24 expression in macrophages leading to lung eosinophilia. Cell symbols were taken from http://smart.servier.com website.
